# Environmental DNA survey captures patterns of fish and invertebrate diversity across a tropical seascape

**DOI:** 10.1101/797712

**Authors:** Bryan N. Nguyen, Elaine W. Shen, Janina Seemann, Adrienne M.S. Correa, James L. O’Donnell, Andrew H. Altieri, Nancy Knowlton, Keith A. Crandall, Scott P. Egan, W. Owen McMillan, Matthieu Leray

## Abstract

Accurate, rapid, and comprehensive biodiversity assessments are critical for investigating ecological processes and supporting conservation efforts. Environmental DNA (eDNA) surveys show promise as a way to effectively characterize fine-scale patterns of community composition, but most studies to date have evaluated its effectiveness in single habitats and for conspicuous taxonomic groups in temperate ecosystems. We tested whether a single PCR survey of eDNA in seawater using a broad metazoan primer could identify differences in community composition between five adjacent habitats at 19 sites across a tropical Caribbean bay in Panama. We paired this effort with visual fish surveys to compare methods for a conspicuous taxonomic group. eDNA revealed a tremendous diversity of animals (8,586 operational taxonomic units), including many small taxa that would be undetected in traditional in situ surveys. Fish comprised only 0.07% of the taxa detected by a broad COI primer, yet included 43 species not observed in the visual survey. eDNA revealed significant differences in fish and invertebrate community composition across adjacent habitats and areas of the bay driven in part by taxa known to be habitat-specialists or tolerant to wave action. Our results demonstrate the ability of broad eDNA surveys to identify biodiversity patterns in the ocean.

## Introduction

Coastal regions make up less than 10% of the Earth’s surface but their ecosystems contribute disproportionately to the globe’s primary productivity, biodiversity, and ecosystem services ^1–4^. Coral reefs alone are thought to be home to 25% or more of described marine species^5^. Human activities such as coastal development, exploitative fishing practices, and eutrophication, however, have resulted in the widespread decline of commercially-important fisheries, the loss of important habitat-forming species, and biological invasions^6–8^. The accelerating pace of changes in the structure and function of coastal ecosystems due to these impacts makes it urgently important to develop methods for efficient and effective biomonitoring to support management, conservation, and basic science initiatives^9^.

Systematic survey data are especially fundamental in understanding the link between biodiversity and the health and functioning of marine ecosystems^10–12^. At its core, biomonitoring requires the reliable identification of species that are present in an environment, and answers if, how, and why populations of these species change over time. Yet, it still remains a challenge to capture the full taxonomic diversity of ecosystems in a repeatable way to identify trends through time and patterns across space^13^. Traditional marine biodiversity surveying methods, such as visual surveys by divers, are often expensive, invasive, require taxonomic expertise, are limited by visibility or habitat complexity, and miss cryptic diversity, including most invertebrates^14–16^. As a result, such traditional techniques are prone to miss many species or even entire taxonomic and functional groups, and are sensitive to observer bias^14,17–20^.

Sampling environmental DNA (eDNA), the genetic material shed by organisms into the surrounding environment (e.g., blood, mucus, waste, scales, etc.^15^), may avoid the shortcomings of traditional survey techniques, and thereby provide a powerful and repeatable approach to assess biodiversity^15,16,21–26^. Studies have shown the promise of eDNA metabarcoding for detecting specific target taxa and elucidating richness and community composition patterns in taxonomically focused surveys using species- or group-specific primers. Relatively few marine studies, however, have explicitly tested whether broad range eDNA surveys potentially targeting all metazoans simultaneously in a single PCR assay could effectively detect fine-scale patterns of community composition in spatially heterogeneous coastal seascapes^22,27,28^. Moreover, none of these studies were conducted in the tropics, nor validated with visual surveys on subsets of the sampled communities. In this era of unprecedented biodiversity loss^8,29^, development of a reliable community-wide approach to assess tropical marine biodiversity across greater spatial and temporal scales would provide a more holistic picture of ecosystem health and functioning by establishing patterns of biodiversity and possible species linkages between habitats.

Here, we test the efficacy of a broad metazoan eDNA metabarcoding survey in tropical marine environments and validate the approach with an established visual survey protocol [i.e., Reef Life Surveys (RLS)^30^] for fish. We examined fish and invertebrate communities in multiple interconnected tropical habitats that have traditionally been challenging to survey comprehensively despite their critical functional roles — e.g., they host considerable biodiversity, serve as important nurseries for juveniles, and sustain high levels of productivity^31–35^. The semi-enclosed Almirante Bay in the Bocas del Toro archipelago on the Caribbean coast of Panama is an ideal system for testing the ability of eDNA to capture fine-scale patterns of marine diversity. It contains all the primary tropical coastal habitats including coral reefs, seagrass meadows, mangrove forests, sandy unvegetated bottoms, and human-made structures, each with distinct communities that are in close proximity, forming a heterogeneous seascape^36,37^. Almirante Bay is also one of the best studied areas in the Caribbean, both taxonomically and ecologically. A large-scale biodiversity survey conducted in 2003 and 2004 across the bay^36^ reported 1183 species of marine invertebrates, 128 species of fish^38^ and helped establish an online database of species reported from the area, with photos (available at https://biogeodb.stri.si.edu/bocas_database/). Since then, annual taxonomic workshops have been held in the area^36,39,40^ and the database has expanded to include over 6,500 terrestrial and marine plant, animal, and fungi species. Over 80% of the fishes reported by the database are represented in GenBank for the animal barcode cytochrome c. oxidase subunit I (COI) gene. Almirante Bay has also served as a natural laboratory for numerous ecological studies utilizing visual survey and experimental methods to look at the response of coral reefs and associated reef fauna to anthropogenic and environmental stress^41–45^.

The objectives of our study were to 1) examine patterns of community composition for metazoans using eDNA and 2) validate the resolution of a broad COI metazoan primer by comparing the subset of fish taxa detected by eDNA to those identified in traditional visual fish surveys and previously assembled taxonomic lists of reported species. We hypothesized that despite the potential for eDNA in seawater to mix across habitat boundaries^27^, eDNA surveys would detect habitat-specific patterns of community composition and structure across the highly diverse mangrove-seagrass-coral reef complex.

## Materials and Methods

### Study Sites

The field component for this study was conducted in Almirante Bay of the Bocas del Toro archipelago (Figure 1) on the Caribbean coast of Panama from May to July 2017. A total of 19 study sites that span a wide gradient of environmental conditions were sampled; they were separated by distances ranging from 20 m to 6,000 m and included areas exposed to, and sheltered from, open ocean swell (Figure 1, Table S1). Six of these areas were previously established Smithsonian Marine Global Earth Observatory (MarineGEO) monitoring sites that each contained adjacent mangrove, seagrass, coral reef and sandy bottom habitats. Eight additional sites contained only mangrove and seagrass habitats, and five were boat docks (Figure 1). Additional site information can be found in Supplementary Table S1. Surveys in mangrove habitat were conducted in and around the submerged portion of *Rhizophora mangle* stands (1-2 m depth). Surveys in seagrass habitat were conducted in meadows dominated by *Thalassia testudinum* (2-4 m depth). Surveys in coral habitat were conducted in reef areas that were dominated by *Porites* spp. finger corals and *Agaricia tenuifolia* (2-5 m).

**Figure 1.**
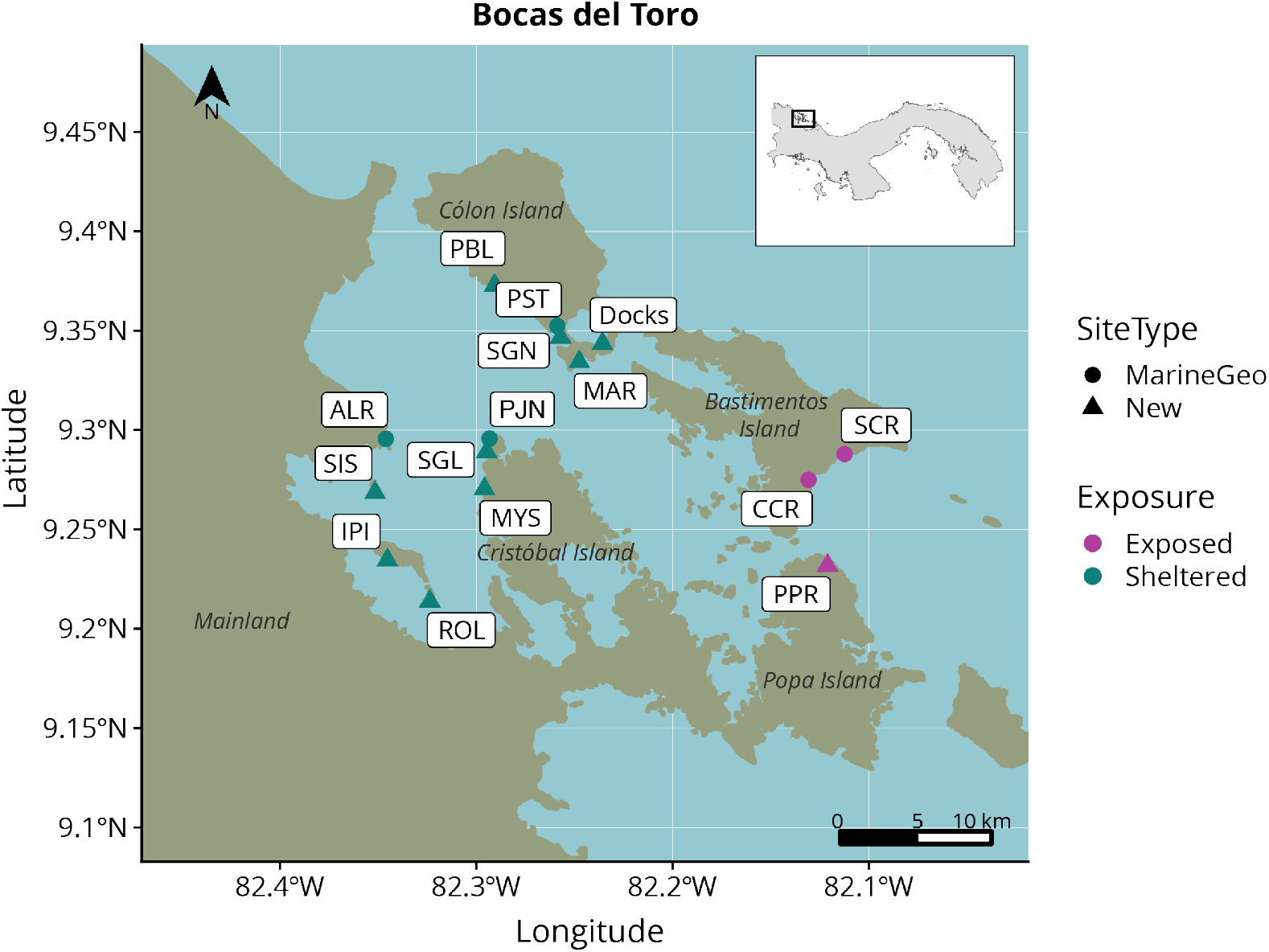
Sampling sites. Samples were collected in MarineGEO sites (circles) from mangrove, seagrass, coral, and sand habitats, whereas additional sites (triangles) were only sampled in mangrove and seagrass habitats, or at boat docks. Full site names can be found in Supplementary Table S1.

### eDNA Surveys

The eDNA workflow involved filtering and extracting eDNA from a water sample and amplifying specific regions of DNA using a broad COI metazoan primer set during PCR. The resulting amplicons were then sequenced on a massively parallel sequencing platform and compared to a reference database of partially or fully characterized genomes of organisms to identify the taxa present in the environmental sample^26^.

#### Sample collection

Three replicate one-liter seawater samples were collected from each of four habitat types (mangrove, seagrass, coral, sand) at the six MarineGEO monitoring sites (n = 72) and from mangrove and seagrass habitats at eight additional sites (n = 48). At the five dock sites, a 1 L water sample was taken from each of the two sides of each dock; there was one dock at three of the sites and 2 docks at two of the sites (n = 14). In total, 134 water samples were collected (see Supplementary Table S1).

Water samples were collected by opening a translucent bleach-sterilized wide-mouth polypropylene sampling bottle directly above (10 – 30 cm) the surface of the habitat without disturbing benthic sediments. Samples were filtered immediately on the boat after collection when possible, but occasionally frozen at −20°C until filtering. There was no difference in the amount of DNA extracted (t-test, P = 0.55) or in PCR yield (P = 0.14) from freshly filtered versus frozen samples (Supplementary Figures S1 and S2). Water samples were vacuum-filtered onto sterile 47 mm diameter 0.45 μm cellulose nitrate filters (GE Life Sciences, Pittsburgh, PA and ThermoScientific, Rochester, USA) with a peristaltic pump head following the “Environmental DNA Sampling Protocol #2” prepared by the U.S. Geological Survey^46^. One 1 L field blank was also collected per sampling event (n = 19 total) as a negative control (a 1 L bottle of deionized water was left open during filtration of the triplicate eDNA samples in the field and then filtered as described above). Filters were stored in sterile 15 mL falcon tubes at −20°C, then transferred to a −80°C freezer before extraction.

#### DNA Extraction

Half of each cellulose nitrate filter was cut up into smaller pieces for DNA extraction using sterile snips and the other half was archived at −80°C. DNA was extracted using the DNeasy PowerSoil Kit (Qiagen, Carlsbad, CA) with the following modifications to the manufacturer’s protocol. First, we increased the total volume of Powerbead solution (950μL total) and C1 solution (80 μL total) to fully submerge filter pieces. Second, we incubated samples in a water bath at 65°C for 10 minutes directly following bead beating to increase the yield. Third, we eluted the DNA in 75 μL of Solution C6 to increase the final concentration. The concentration of each DNA extract was quantified using the Qubit dsDNA HS Assay (Invitrogen, Ca, USA), and the approximate molecular weight of a selected number of DNA extracts was visualized on a 1.5% agarose gel by electrophoresis.

#### Amplification and Sequencing

A 313 bp fragment of the variable region of the mitochondrial COI gene^47^ was amplified for each sample with a set of PCR primers designed to target metazoans (mlCOIintF and jgHCO2198, full primer sequences can be found in Supplementary Table S2). To combine samples into a sequencing run on the Illumina MiSeq platform, each sample was assigned a unique combination of indices following the adapter ligation method^48^. The first index was added to the target fragments during PCR amplification using tailed-PCR primers (i.e., PCR primers to which unique 6-base-pair indices were added at the 5’ end) and the second index was added to the product of amplification in the form of standard single-indexed Illumina TruSeq Y-adapters. PCR amplifications were conducted three times for each sample. Each 20 μL PCR reaction contained 2 μL of Clontech Advantage 2 PCR buffer (10x), 1 μL of each primer (10μM), 1.4 μL of dNTP mix (10 mM), 0.4 μL of 50X Advantage 2 Polymerase Mix (50x), 13.2 μL PCR-grade water, and 1 μL of DNA extract. The following cycling conditions were used: 5 minutes at 95°C (1x); 1 min at 95°C, 30 s at 48°C, and 45 s at 72°C (38x); 5 min at 72°C (1x). PCR negative controls were conducted alongside samples to test for the presence of contaminants. PCR replicates were pooled, purified using magnetic KAPA Pure Beads (bead:sample ratio of 1.6:1) and quantified using Qubit dsDNA HS Assay. Equimolar amounts of PCR products amplified with distinct tailed PCR primers were pooled before ligation of single-indexed adapters (see Supplementary Table S3 for the multiplexing protocol) following the protocol of the Illumina TruSeq PCR-Free LT kit. All libraries were quantified using a Qubit fluorometer, equimolar amounts pooled into a single tube, and the final product validated using a KAPA qPCR library quantification kit (KAPA Biosystems, Wilmington, Massachusetts, USA). The library was sequenced on an Illumina MiSeq with an Illumina MiSeq v2 500-cycle kit.

We also attempted to amplify and sequence the water samples using the fish-specific 12S MiFish primers for additional validation, but we had limited success with PCR amplification using these primers. This may be due to the low concentration of fish DNA present in our water samples. For this reason, we do not report further on the fish-specific amplicon sequencing part of this study, but additional details for the MiFish amplicon method used can be found in the Supplementary Information document.

### Analysis of sequence data

Reads were demultiplexed and Illumina adapters were trimmed using Flexbar version 3.0.3^49^. The demultiplexed and adapter trimmed FASTQ files can be accessed on the NCBI SRA under the BioProject accession number PRJNA558350. Negative control samples that failed DNA extraction, PCR amplification, or initial quality control checks after demultiplexing were not uploaded. Adapter-trimmed reads were then further trimmed to ensure that all primer regions were removed, filtered for quality, merged, checked for chimeric sequences, and used to infer exact amplicon sequence variants (ASVs) with DADA2 version 1.9.0^50,51^ (DADA2 parameters: maxN = 0, maxEE = c(2,2), truncQ = 10, trimLeft = 26, with pseudo-pooling). ASVs were clustered into operational taxonomic units (OTUs) at a 97% identity threshold using VSEARCH^52^. OTUs were further processed with the LULU algorithm (GitHub commit f4a880d), which curates OTUs based on sequence identity and co-occurrence patterns, in order to reduce taxonomic redundancy and improve the accuracy of richness estimates (LULU parameters: minimum_ratio_type = “min”, minimum_ratio = 1, minimum_match = 84, minimum_relative_cooccurence = 0.95)^53^. OTUs were assigned taxonomic information using the Bayesian Least Common Ancestor (BLCA) Taxonomic Classification software^54^ (GitHub commit 3b41b98) against Midori-Unique v20180221, a curated database of metazoan mitochondrial gene sequences (available at www.reference-midori.info)^55^. BLCA taxonomic rank assignments with less than 50% confidence were discarded. OTUs that remained unidentified with BLCA were compared to the whole NCBI NT database (retrieved May 2018) using BLAST searches (word size = 7; max e-value = 5e-13) and assigned the taxonomic information of the lowest common ancestor of the top 100 hits.

OTUs that were identified down to species level were cross-checked with the species lists from the Smithsonian Tropical Research Institute’s Bocas del Toro species database (https://biogeodb.stri.si.edu/bocas_database/, retrieved 3/9/2019), previous literature on Caribbean biodiversity^13^, the Ocean Biogeographic Information System for the Bocas region (OBIS, retrieved 6/2/2019), and the Global Invasive Species Database (retrieved 4/11/2019). All R code and input files can be found on GitHub at: https://github.com/Talitrus/bocas_eDNA.

### Visual Fish Surveys

To quantify the richness and composition of fish communities in coastal habitats across the network of 19 sites, two scuba divers conducted visual fish surveys following the Reef Life Survey (RLS) protocol^30^. These divers counted, identified to the highest taxonomic resolution possible, and assigned to binned size-classes all fish greater than 2.5 cm in length observed along a 50 × 10 m belt transect positioned parallel to shore. One transect was conducted in each of the same habitats and sites as the eDNA survey (Figure 1, Table S1) for a total of 43 transects. Due to the inability to swim through the mangrove habitat, two 50 m transect lines were deployed along the edge of the mangrove and a diver looked 5 m to one side of each of the primary prop roots and under overhanging roots for fish. These mangrove transects were then pooled together into a single transect. Dock sites were sampled similarly to the mangroves, with divers swimming along the edge of the structure looking inward. When fish were in large schools, counts were estimated to the nearest power of ten. Unidentifiable individuals were photographed for later identification using reef fish identification books^56^ and online databases (e.g., FishBase^57^, Smithsonian Identification Guide – Shorefish of the Greater Caribbean^58^). The same two divers conducted all visual surveys after a period of training to ensure consistency.

### Analysis of diversity patterns

Community analyses were conducted in R version 3.5.1^59^ using the *phyloseq*^60^, *vegan*^61^ and *DESeq2*^62^ packages. Machine learning analysis was conducted using the *h2o* package^63^. Visualizations were done with the *Plotly*^64^ and *ggplot2* packages^65^.

Raw counts were transformed to relative abundances before calculating a dissimilarity matrix based on the Bray-Curtis metric. Bray-Curtis takes into account differences in abundance of reads between samples; a value of 0 indicates that samples are exactly identical in terms of OTU composition and relative abundance of reads whereas a value of 1 indicates that samples do not have any OTUs in common. The variance in community composition was calculated using a PERmutational Multivariate ANalysis Of VAriance (PERMANOVA)^66^ computed with 10000 free permutations. The following terms were used as factors: region within the bay, habitat, geographic site, diver (for the visual survey), and the interaction terms between bay region and habitat, and site and habitat.

We calculated the accuracy with which a machine learning algorithm was able to assign a sample to its source habitat based on OTU composition to understand how consistently distinct the assemblage was for a given habitat in Bocas del Toro. We selected a random forest classifier over other approaches because of the high-dimensionality of our data, the relative ease of tuning a distributed random forest algorithm, and the ability of random forest algorithms to calculate feature importance. LULU-curated metazoan OTUs were filtered to only include taxa that occurred at more than 5e-5 relative abundance (approximately five reads on average) in at least five samples. OTUs were then used as predictors in a random forest classifier with source habitat (coral reef, mangrove, seagrass, sand, dock) as the response variable with 5-fold cross-validation using the h2o machine learning platform^63^. Cross-validation is a statistical term for training a model on the majority of the data while withholding some data to be used to test the model. Since the withheld data were not used to train the model, the model has never seen them before, thereby providing a better estimate of how accurate the model would be on new data. Hyperparameters were chosen by grid search optimizing for logarithmic loss using ‘h2o’. The model parameters with the smallest logarithmic loss value during cross-validation were selected and used to build a random forest from which feature importance information was extracted.

To examine differences in specific pairwise habitat and exposure comparisons (exposed vs. sheltered, mangrove vs. seagrass habitat, and coral vs. sand habitat), differential abundance analysis of visual survey detections and OTUs of fish and metazoans between habitats was conducted using Wald t-tests in the DESeq2 package^62^. Coral and sand sites were only contrasted with each other because each only had six sites, while mangrove and seagrass sites had 12 sites each. Variance stabilizing transformation was used on the count data to standardize downstream visualization using heatmaps.

## Results

### Metazoan community composition

PCR amplification of the mitochondrial COI gene with a general primer set was successful for all 134 water samples whereas all negative field, extraction and PCR controls remained blank. A total of 14,376,785 raw paired end reads were obtained and 12,708,398 (88.4%) remained after filtering, chimera removal, and processing.

The vast majority of the reads (99.5%) from seawater were assigned to non-fish taxa such as invertebrate animals, unicellular eukaryotes (i.e., diatoms and picozoans), algae or bacteria (Figure 2). Metazoans made up 8,781 OTUs (out of 23,123 total OTUs) consisting of 5,479,007 paired reads (43% of total). Of these, approximately 10% of sequences were identified to species (526,898 read pairs comprising 387 OTUs). OTU richness and alpha diversity (Shannon’s Index, H’) was highest in mangrove habitats (4,939 OTUs, 5.127 ± 1.63) and lowest in coral reef habitats (1,680 OTUs, 4.912 ± 0.069) (Table 1).

**Figure 2.**
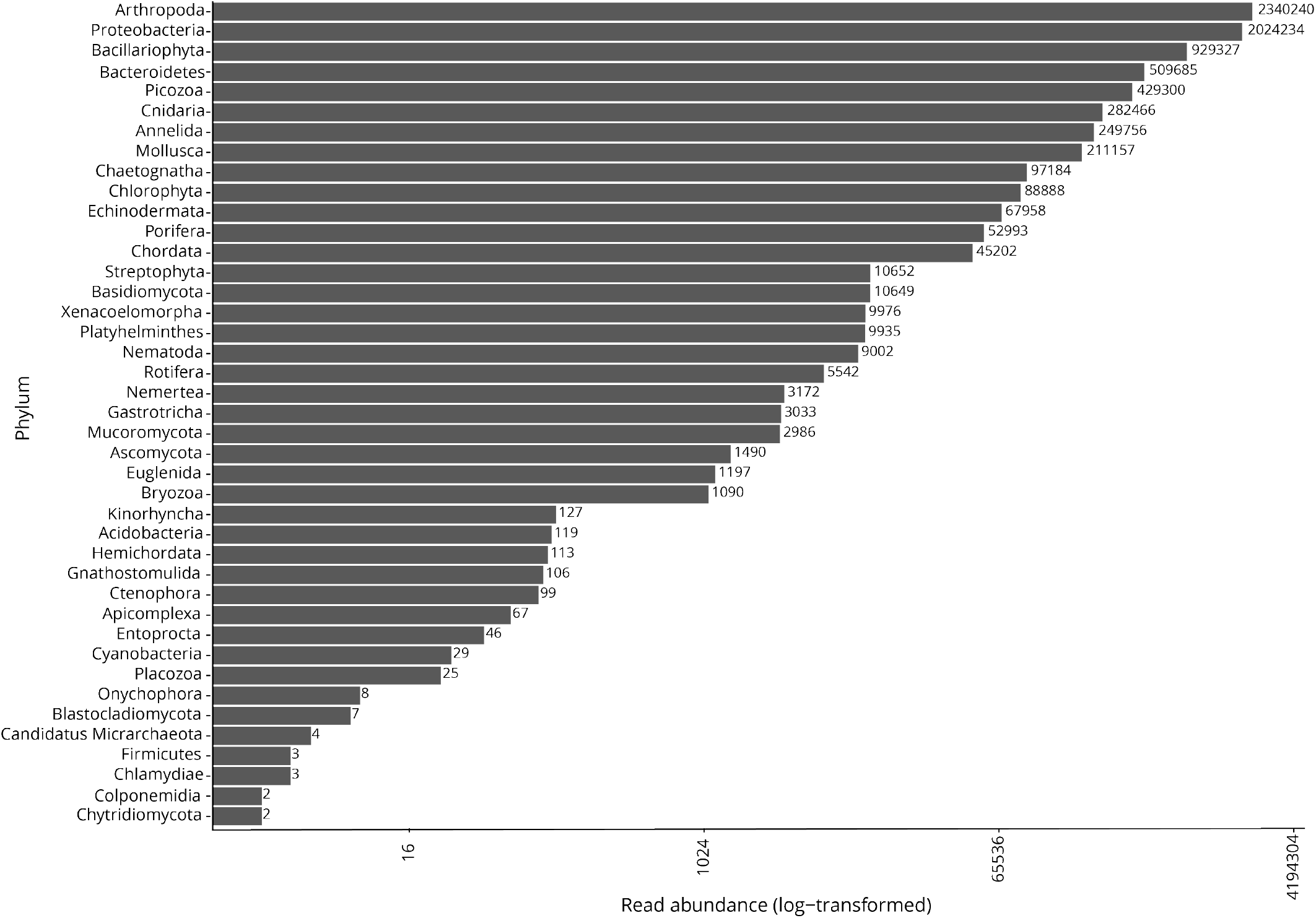
Total read abundance for phyla detected after filtering and quality controlling eDNA samples at all 19 sites sampled.

**Table 1.**
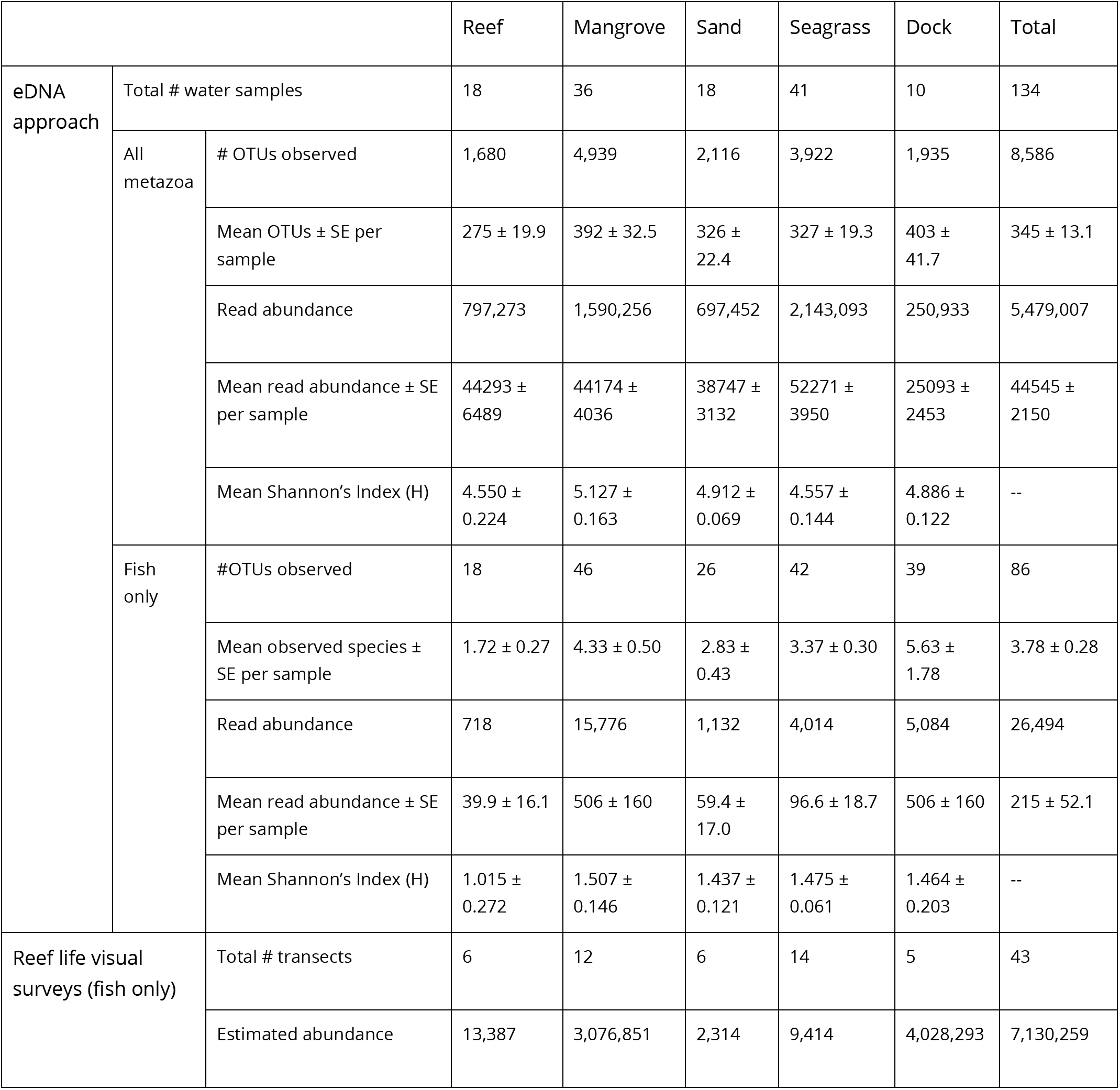
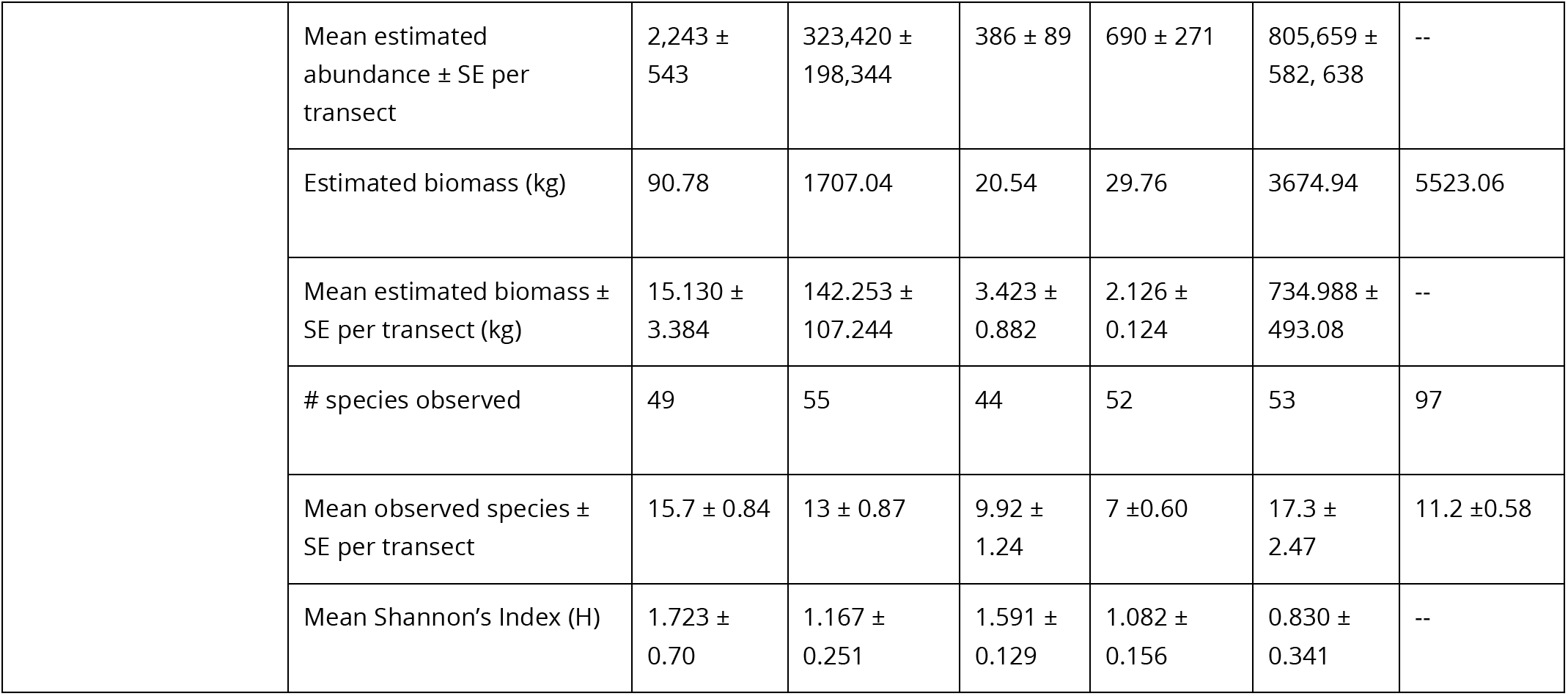
Summary table of taxonomic richness, diversity, and abundance detected by eDNA and visual surveys at all sites.

A larger portion of the variation in the metazoan eDNA data was explained by sampling site (R^2^=29.1%) and the interaction term between habitat and sampling site (R^2^=17.2%). PERMANOVA analysis indicated significant differences between geographic areas (exposure), habitats, sites and all interactions between factors (Table 2). Despite targeting a much broader range of taxa (including fish and numerous invertebrates), the whole metazoan eDNA sequencing found similar patterns of community dissimilarity across the sampling region and between habitats (PERMANOVA, Table 2) as the visual and the subset of the eDNA dataset assigned to fish.

**Table 2.** PERMANOVA results using Bray-Curtis distances for (a) eDNA metazoan surveys, (b) eDNA fish surveys and (c) Reef life visual fish surveys testing for differences in community composition. Asterisks represent p-values equal to or smaller than: 0.05 (*), 0.01 (**), and 0.001 (***).

| (a) | Term | Df | SS | F Model | R<sup>2</sup> | P-value |  |
| --- | --- | --- | --- | --- | --- | --- | --- |
| eDNA all metazoa | Exposure | 1 | 1.440 | 7.436 | 0.033 | 0.001 | *** |
|  | Habitat | 4 | 4.222 | 5.452 | 0.098 | 0.001 | *** |
|  | Site | 15 | 12.505 | 4.306 | 0.291 | 0.001 | *** |
|  | BayRegion: Habitat | 3 | 1.966 | 3.385 | 0.046 | 0.001 | *** |
|  | Habitat: Site | 19 | 7.380 | 2.001 | 0.172 | 0.001 | *** |
|  | Residual | 80 | 15.488 |  | 0.360 |  |  |
|  | Total | 122 | 43.000 |  | 1.000 |  |  |

| (b) | Term | Df | SS | F Model | R<sup>2</sup> | P-value |  |
| --- | --- | --- | --- | --- | --- | --- | --- |
| eDNA fish | Exposure | 1 | 1.368 | 4.000 | 0.026 | 0.001 | *** |
|  | Habitat | 4 | 4.084 | 2.985 | 0.078 | 0.001 | *** |
|  | Site | 15 | 8.754 | 1.706 | 0.168 | 0.001 | *** |
|  | Exposure:<br>Habitat | 3 | 1.760 | 1.715 | 0.034 | 0.001 | *** |
|  | Habitat:Site | 19 | 10.759 | 1.655 | 0.207 | 0.001 | *** |
|  | Residual | 74 | 25.313 |  | 0.486 |  |  |
|  | Total | 116 | 52.038 |  | 1.000 |  |  |

| (c) | Term | Df | SS | F Model | R<sup>2</sup> | P-value |  |
| --- | --- | --- | --- | --- | --- | --- | --- |
| Visual fish | Exposure | 1 | 0.676 | 2.158 | 0.020 | 0.008 | ** |
|  | Habitat | 4 | 1.737 | 1.386 | 0.051 | 0.030 | * |
|  | Diver | 1 | 0.033 | 1.060 | 0.009 | 0.387 |  |
|  | Site | 15 | 8.594 | 1.829 | 0.255 | 0.001 | *** |
|  | Exposure:<br>Habitat | 3 | 1.874 | 1.994 | 0.056 | 0.001 | *** |
|  | Habitat:<br>Site | 19 | 8.620 | 1.448 | 0.256 | 0.002 | ** |
|  | Residual | 38 | 11.905 |  | 0.353 |  |  |
|  | Total | 81 | 33.737 |  | 1.000 |  |  |

A distributed random forest algorithm correctly predicted the source habitat of the samples an average of 83% and 68% of the time in total and in cross-validation, respectively. Mangrove samples were the most likely to be correctly classified during cross-validation (81% correct) while coral reef samples were the least likely (44%) (Figure 3). While many of the taxa that were important predictors in the random forest classification algorithm were unidentified taxa, a number of identified taxa were species known to occupy mangrove roots, such as *Haliclona manglaris, Isognomon alatus*, and *Dendostrea frons*. (Supplementary File S1).

**Figure 3.**
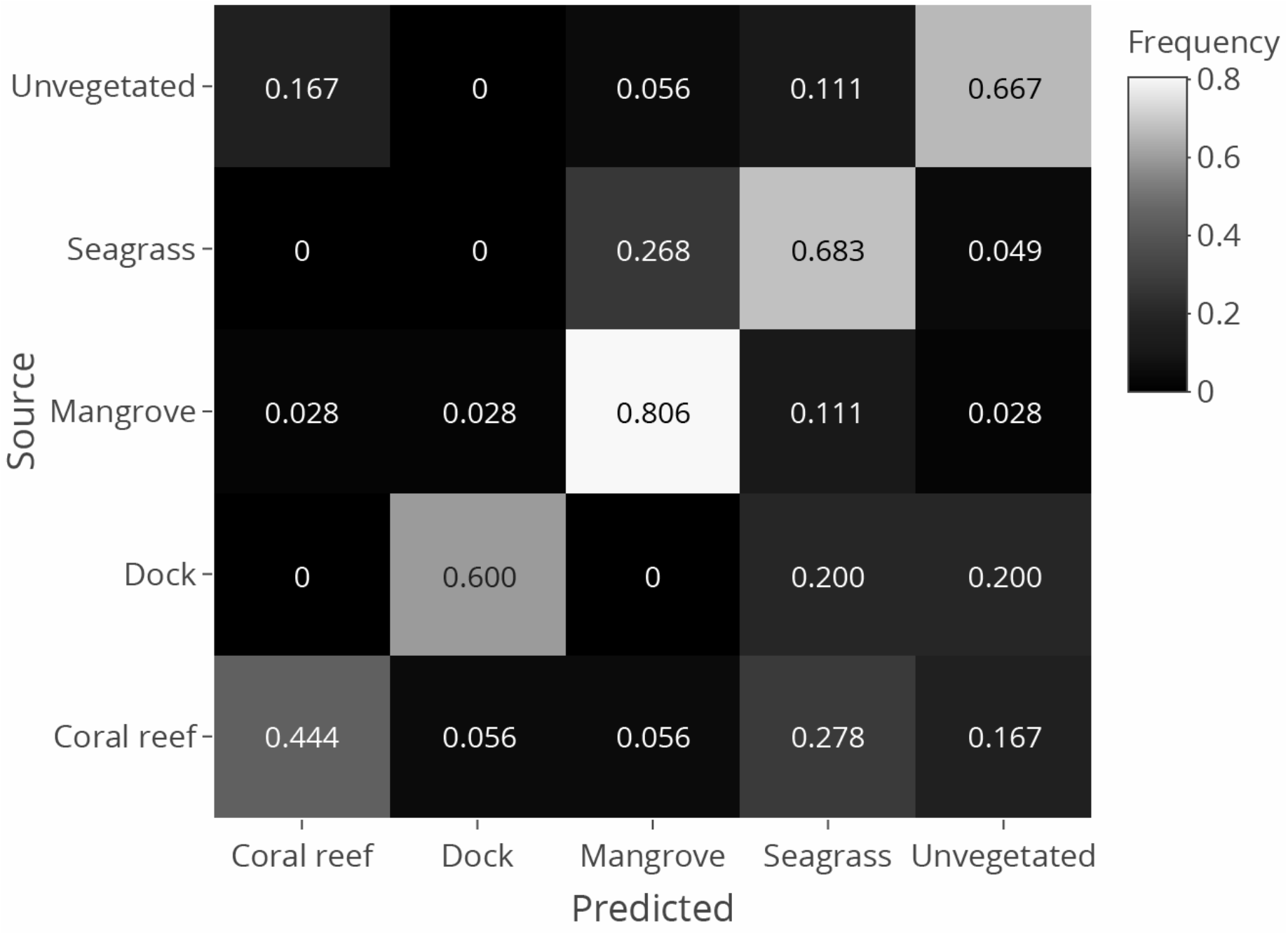
Frequency at which samples from each habitat were classified into predicted habitat types using random forest model. The confusion matrix shows hold-out prediction frequencies during cross-validation. The cells on the counter-diagonal represent correct classifications. The frequency is calculated out of the total samples from each source respectively (i.e., each row adds up to 1, but not necessarily each column).

The identity and abundance of metazoan OTUs differed between mangrove and seagrass habitats (Supplementary Figure S3), between reefs and sand habitats (Supplementary Figure S4), and by exposure (Figure 4). There were more OTUs that significantly (adjusted p-value < 0.05) differed in abundance when comparing exposed versus sheltered sites than any of the habitat comparisons. Samples generally consisted of invertebrate OTUs that were unidentifiable past the phylum level. Of eDNA OTUs with species-level assignment, the symmetrical brain coral, *Pseudodiploria strigosa*, and sponge *Placospongia carinata* were more abundant in exposed sites, while the mangrove periwinkle (*Littoraria angulifera*), European anchovy (*Engraulis encrasicolus*), eared ark clam (*Andara notabilis*) and planktonic copepod *Oncaea waldemari* were more abundant in sheltered sites (Fig 4).

**Figure 4.**
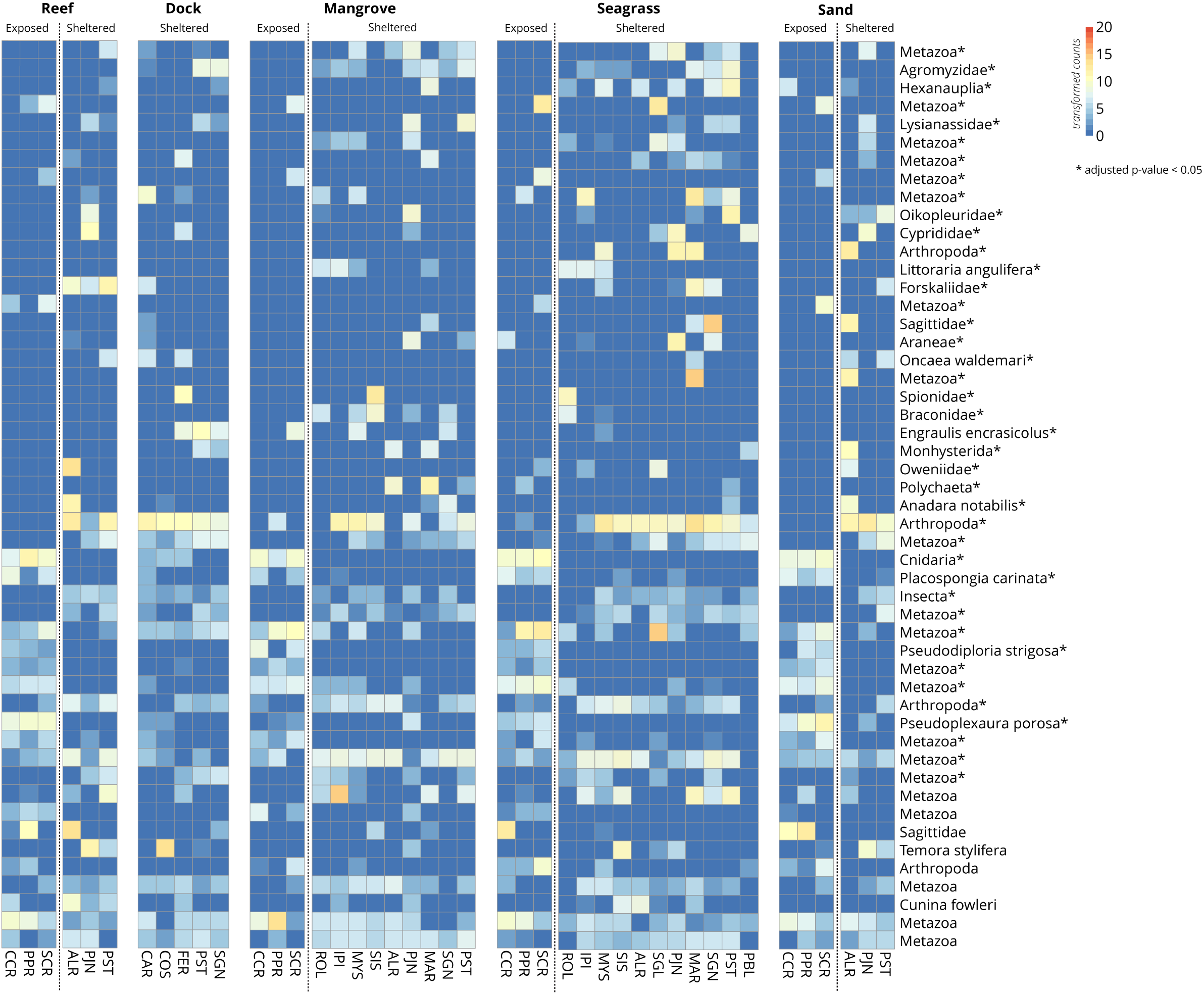
Heatmaps of metazoan taxa differentially abundant in exposed and sheltered sites. The top 50 taxa are ordered top to bottom by ascending adjusted p-values (decreasing significance), with asterisks (*) denoting an adjusted p-value < 0.05. The scale bar and cell colors show count values after variance stabilizing transformation. Taxa labeled as higher than species-level represent unique OTUs that could not be identified below that level. For the full names of these OTUs, refer to Supplementary File S2.

Of the OTUs identified to species level, only 154 (40%) had been previously reported in the Smithsonian Tropical Research Institute Bocas del Toro species database or OBIS at the time of analysis (see Supplementary File S2 for exact lists). Interestingly, four of the species detected in the eDNA, including two terrestrial species, are listed as invasive in the Caribbean by the Global Invasive Species Database: the bryozoan *Bugula neritina*, the crab *Charybdis hellerii*, the ant *Monomorium pharaonis*, and the common house mouse *Mus musculus*. DNA from these invasive species was observed in more than 3 water samples, and all were at very low relative abundances (*Bugula neritina*: 46 reads, *Charybdis hellerii*: 7 reads, *Monomorium pharaonis*: 186 reads, *Mus musculus*: 21 reads). We speculate that the terrestrial invasive species DNA is likely from local sewage sources and/or runoff rather than laboratory contamination, since these are species known to occur in the area and were not found in any negative or positive controls.

### Fish Community Composition

In the eDNA survey, reads assigned to bony fishes (Osteichthyes) made up a median of only 0.07% of the total reads per sample with no sample containing more than 5.2% fish reads. We detected 79 fish species (86 OTUs) which accounted for a total of 26,724 paired reads (Table 1). 93% (80/86 OTUs) had a species-level identification, with two OTUs identified as the same species. 66.6% (4/6) of OTUs only had a family-level identification, and 33% (2/6) of OTUs only had a class-level identification. Six out of 134 eDNA sample replicates did not contain hits to fish, but every habitat within each site had hits to fish when triplicate samples were combined. The highest mean read abundance of bony fishes among habitats was recovered from the dock sites (per sample mean ± SE: 506±160), followed by mangrove habitats (per sample mean ± SE: 436±160) (Table 1).

Mangrove habitats harbored the highest average read diversity for fish based on Shannon diversity indices, followed by seagrass, dock, sand, and finally coral reef habitats (Table 1). Samples from the Salt Creek mangrove MarineGEO site, located outside the bay, contained among the highest number of hits to fish species (10,229 reads), followed by the Ferry dock site located inside the bay (1,510 reads). The fish species with the highest number of reads were the clupeid big-eye anchovy (*Anchoa lamprotaenia*; 11,312 reads), striped parrotfish (*Scarus iseri*; 1,700 reads), and clupeid hardhead silverside (*Atherinomorus stipes*; 1,671 reads). Sheltered sites had a higher species richness and diversity (6034 OTUs, Shannon’s H= 5.172) than exposed sites (4781 OTUs, Shannon’s H = 4.934).

For the visual fish survey, over 7,000,000 individuals (5,523 kg) and 97 identifiable species from 42 families were observed in transects covering a total area of 21,500 m^2^ (Table 1). At least one fish individual was observed from each habitat in every site. Fishes that were ubiquitous across all sites and habitats belonged to species that are typically thought of as reef-associated, were already included in the Bocas species database, and were previously observed in surveys of the area (e.g., stoplight parrotfish (*Sparisoma viride*), threespot damselfish (*Stegastes planifrons*), bluehead wrasse (*Thalassoma bifasciatum*)). Of the non-schooling fish species, the masked goby (*Coryphopterus personatus*) was the most abundant (and third most abundant overall). For a full list of fish species, see Supplementary Table S4.

Out of a total of 922 diver observation events (*i.e*., a species or morphospecies recorded in some abundance on a given transect by a given diver), 838 (91%) were species-level identifications, 20 (2%) were genus-level identifications and 64 were family-level identifications (7%). The family-level observations accounted for 99% of the observed individuals across all study sites. Clupeids (small mid-water “baitfish”) commonly found in mangrove and dock habitats could not be identified to species in most cases because they are small, move quickly, and occur in very large mixed species schools (e.g., up to 1,000,000 estimated in a single transect). Clupeids and the second most abundant group, belonids, accounted for 75% of diver observation events that were not identified at the species level. For fishes other than clupeids and belonids, 98% of diver observation events achieved species-level identifications.

### Comparing fish eDNA and visual survey methods

In both the visual surveys and eDNA surveys, we found significant differences in the fish communities observed between different regions of the bay, and between habitats, using the Bray-Curtis index based PERMANOVA (full results in Table 2). There was no clearly visible structure when the community data from the eDNA or visual surveys were ordinated using a Bray-Curtis PCoA (Figure 5). For both surveys, the majority of the explained variation, however, was attributed to the factor “site” which encompasses multiple habitats within a given location, (RLS: R^2^=25.5%, eDNA: R^2^=16.8%) and the interaction term between habitat and site (RLS: R^2^=25.6%, eDNA: R^2^=20.7%).

**Figure 5.**
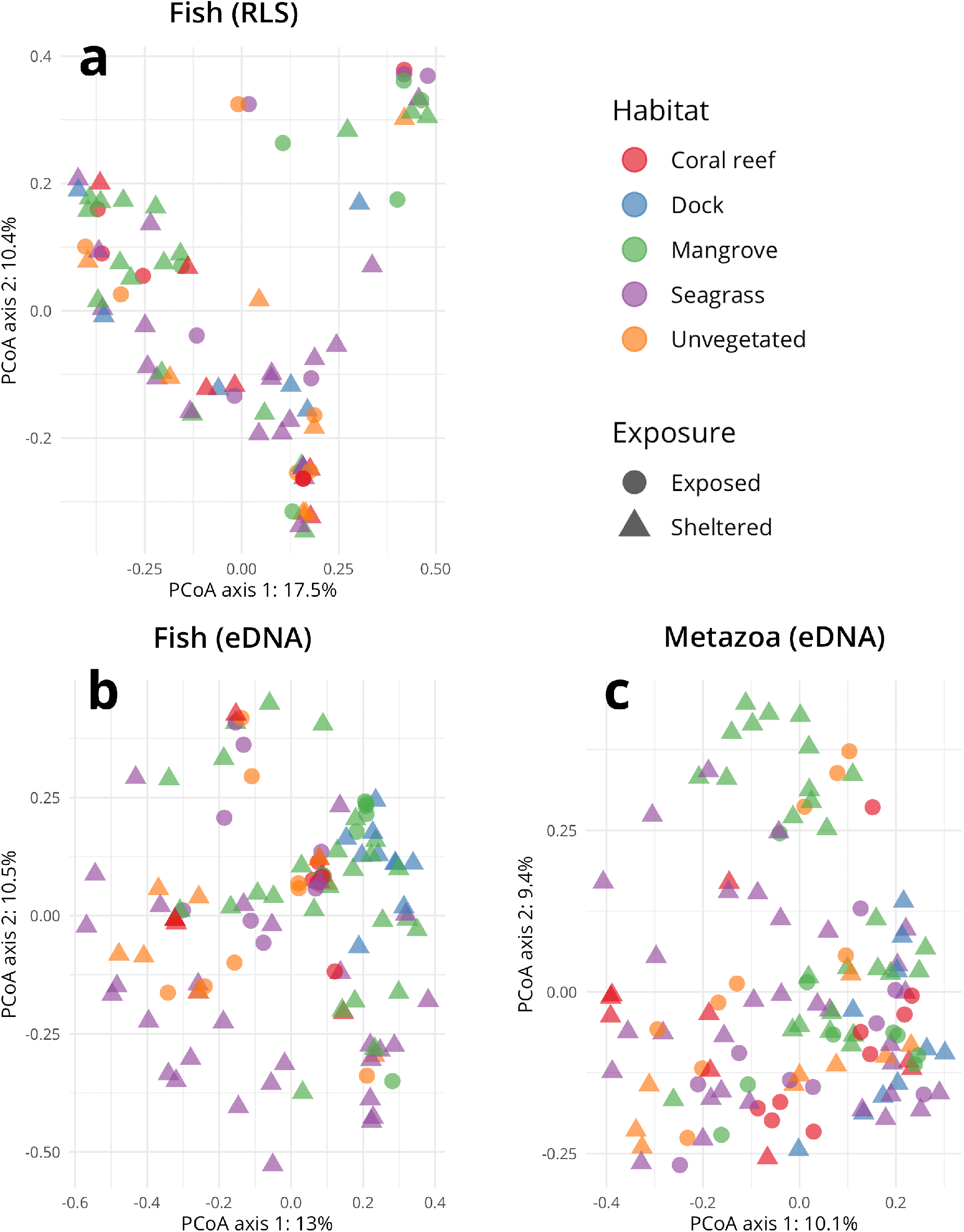
Principal coordinate ordinations for the (a) Reef Life visual fish survey data, (b) fish detected in eDNA with COI primers, and (c) all metazoans detected in eDNA with COI primers. Percentages in the axis titles represent the proportion of that axis’ eigenvalue to the sum of all eigenvalues.

Visual surveys and eDNA were similar in that each method identified differences between habitats and exposure in the abundance of fish taxa, but they differed in which taxa contributed to this difference (Figure 6 and Supplementary Figure S4). For example, in visual surveys, nine taxa including schoolmaster snapper (*Lutjanus apodus*), latin grunt (*Haemulon steindachneri*), and yellowfin mojarra (*Gerres cinereus*) were more abundant in mangrove sites, whereas slippery dick (*Halichoeres bivittatus*) was more abundant in seagrass sites (Fig 6). On the other hand, the same comparison between mangrove and seagrass sites for the eDNA fish dataset only showed big-eye anchovy (*Anchoa lamprotaenia*) to be more abundant in mangrove sites (Figure 6). In general, the visual surveys detected more differentially abundant taxa in the habitat and wave exposure comparisons than the eDNA surveys.

**Figure 6.**
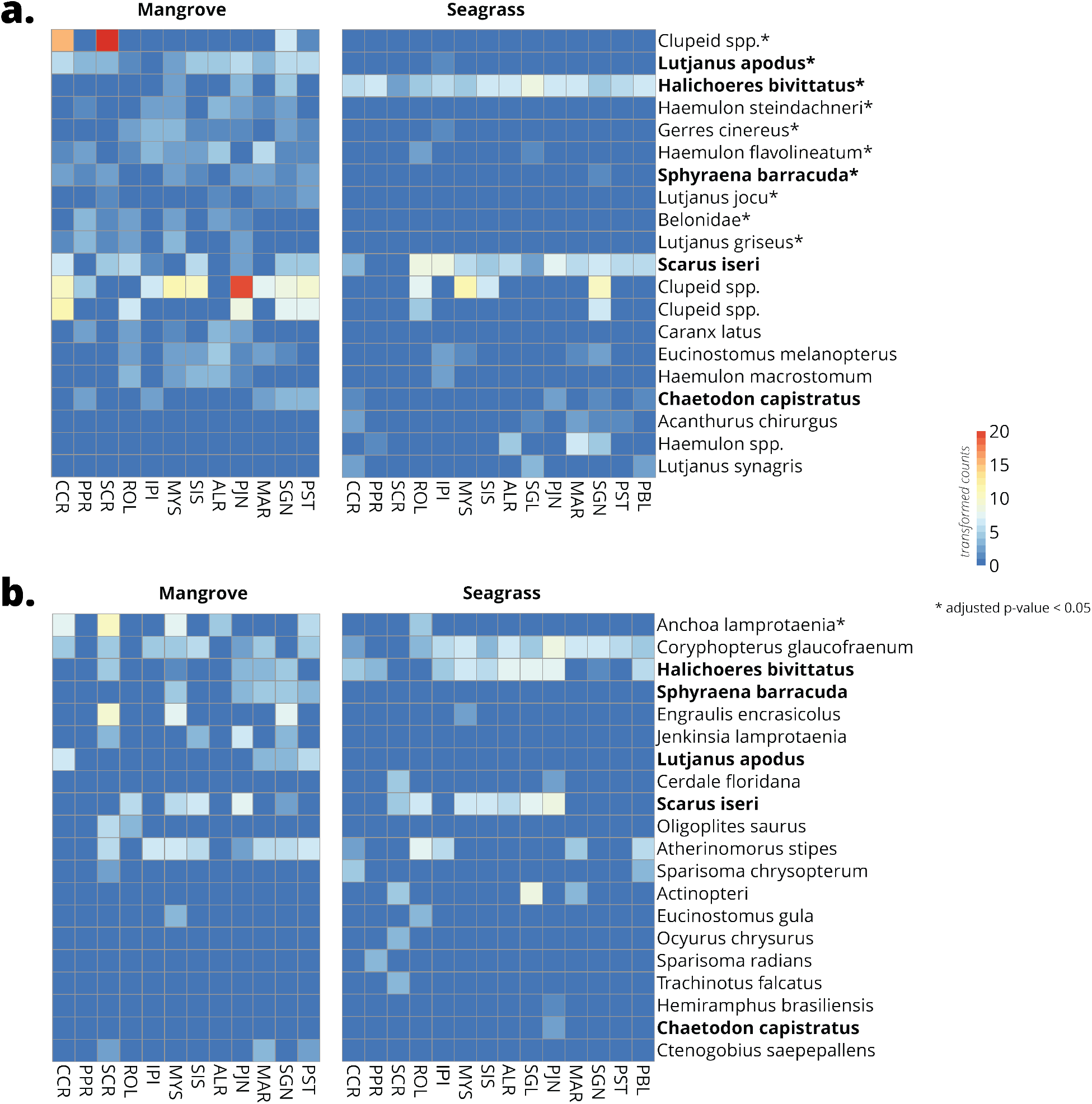
Heatmaps of the top 20 bony fish taxa differentially detected in mangrove and seagrass habitats using (a) Reef Life visual fish surveys and (b) eDNA surveys. The 20 fish taxa are ordered from top to bottom by ascending adjusted p-values (decreasing significance), with asterisks (*) denoting an adjusted p-value < 0.05. Bolded taxa are shared between visual and eDNA surveys. The scale bar and cell colors show count values after variance stabilizing transformation.

The visual survey detected 97 fish species, 36 of which overlapped with the eDNA survey and 19 of which were absent in the eDNA (Supplementary Table S4). Of the 79 fish species detected by eDNA, 31 had not been previously recorded in the Bocas or OBIS databases. No detected fish species (except those mis-assigned, see Discussion) were identified as false positives. In total, eDNA metabarcoding with COI and visual surveys detected 140 fish species, 60 of which had not been previously reported in the Bocas database (which had 219 fish species when we initiated our study) and nine of which had not been reported in OBIS. Our total is slightly greater than the number of detected fish species from the most recent and comprehensive record from this region (2005), which reported 128 fish species (85 of which we also report here) from 12 coral reef sites varying in habitat complexity and exposure in Bocas del Toro after 288 visual surveys on comparable transects^38^. With surveys conducted along 43 transects deployed across five habitat types and using two sampling approaches (i.e., visual and eDNA), we found 65 species that were not reported in 2005^38^ and 25 species that were not recorded in the Bocas database. eDNA detected small-sized species (<2.5 cm) and pelagic species that scuba divers did not look for.

## Discussion

The broad-range COI primer set revealed community patterns across a heterogeneous tropical coastal system that were generally consistent with visual surveys of conspicuous fish and the known distribution of invertebrates. Both the eDNA and visual surveys show distinct variation in taxon abundances between habitats and regions in the bay (Figure 4,6, Table 1). Interestingly, the similar patterns of fish community composition uncovered by the two methods were driven by different pools of fish species. This suggests that eDNA can unveil distribution patterns of fish taxa (as well as invertebrates) previously missed or underreported by visual methods. Our results also indicate that habitat and wave exposure drive the distribution of diversity across various taxonomic groups.

Patterns of differential sequence abundance observed between habitats and regions of the bay for metazoan taxa can be verified and explained by their ecology. For example, the massive brain coral *Pseudodiploria strigosa* was consistently detected by eDNA at exposed sites, as would be expected because of their wave resistant morphology^67^. This result is also consistent with previous descriptive studies of Bocas del Toro’s reefs, where this species was found more often at sites on the exposed eastern side of Bastimentos Island than on sheltered sites around Colón Island^68–70^.

The proportion of the variance explained by site (which encompass multiple habitats) in the PERMANOVA (visual: R^2^ = 0.238, eDNA: R^2^ = 0.270, Table 2) may be indicative of a shared species pool and/or connectivity of habitats in the ecosystem (i.e., the mangrove-seagrass-coral complex) through organism dispersal, migration or movement of seawater that carries eDNA. The variance explained by habitat alone was statistically significant, but relatively small compared to site and site × habitat, indicating differences between habitats at some sites but not others. The physical distance between water samples may have been a larger contributor to the community similarity than the habitat of collection. Alternatively, this could be due to daily differences in weather patterns if wind-driven water movement transported eDNA across habitats at some sites (i.e., as collections were made at all habitats of a site in the same day).

The random forest classification was able to detect subtle variation that was informative for classifying the source habitat of the samples, even though only approximately 7% of the variance in Bray-Curtis distances was explained by habitat. Most of the taxa that were important predictors of habitat classification were metazoans not identified to species, but the taxa that could be identified to species tended to be taxa that are known to be specialized on particular habitats. For instance, a number of common mangrove root epibionts were important to the algorithm’s classification accuracy, including the sponge *Haliclona manglaris* and various bivalves such as the flat tree oyster *Isognomon alatus* and the mangrove oyster *Crassostrea rhizophorae*. Because mangroves had the most taxa, they likely contained more unique species associated with it, which improved the accuracy of the classifier. This shows that eDNA is able to pick up multiple species indicative of different habitats separated by only a few meters.

This study demonstrates the utility of eDNA as a biomonitoring tool for highly diverse tropical seascapes. In combination with a traditional visual survey, eDNA allowed for a more comprehensive survey of the biodiversity contained in the Bocas del Toro lagoon. Overall, over 8,500 metazoan OTUs were detected in the eDNA, which is comparable to previous estimates of marine metazoan diversity for the entire Caribbean region (10,676 animal species reported in a 2010 study of georeferenced species records and taxonomic lists^13^). We detected over 200 metazoan OTUs with species-level identifications that had not been previously recorded in the Bocas or OBIS databases. This is particularly notable given that the majority (95%) of metazoan OTUs detected in our study were not identifiable to species with the databases accessed at time of analysis. Even though COI is one of the most commonly sequenced genes for animals, with over 4.5 million sequences in the Barcode of Life Data system^71,72^, we speculate that many small or cryptic taxa that are not well taxonomically described are not adequately represented in GenBank, a point highlighted by the Open Tree of Life project^73^. In addition, the broad-range COI primers also detected numerous non-metazoan sequences. These sequences represented 53% of the reads, which is around 2–3 times greater in relative abundance than in previous studies sequencing bulk benthic samples utilizing the same COI primers^74,75^ but comparable to recent estimates for water samples^76^. This is likely attributable to the lower concentrations of metazoan eDNA in seawater and the broad taxon compatibility of the COI primer set.

Successful taxonomic identification in eDNA-based surveys rely on the taxonomic coverage of reference databases. Yet, as highly diverse tropical systems remain poorly sampled, the majority of OTUs remained unidentified to the species level. In addition, OTUs in groups known to have slow rates of COI evolution^77^ were incorrectly assigned to species that are not known to occur in the area. For example, one octocoral OTU detected as significantly more abundant in exposed sites was assigned as *Muricea fruticosa*, an eastern Pacific gorgonian species. This OTU is more likely to be one of the many Caribbean gorgonian species with identical COI sequences in the PCR target region (e.g., *Pseudoplexaura porosa* and *Eunicea flexuosa*). Two additional false positives (species not known to inhabit the Caribbean) were detected with the taxonomic assignment method used. Ray OTUs were reported as the Eastern Pacific rays *Aetobatus ocellatus* and *Rhinoptera steindachneri*, but are likely the closely-related Western Atlantic species *Aetobatus narinari* and *Rhinoptera brasiliensis* or *R. bonasus*.

The co-amplification of numerous non-metazoan taxa lowered the sampling depth of our target organisms (metzoan) and likely prevented the consistent detection of rare taxa. The taxonomic selectivity of primers, rather than environmental factors such as organismal DNA shedding rates, has been argued to be the main driver behind differences in detection between eDNA and visual approaches^24^. Using more taxon-specific primers could have improved the detection consistency for organisms of interest. PCR yield using MiFish primers that specifically target fish was surprisingly low across samples. It is possible that 1 L water samples did not contain sufficient fish eDNA for consistent amplification with these primers, possibly owing to the overall low fish biomass in our study area relative to other parts of the Caribbean^78^. Increasing the number of replicates and the volume of water filtered, along with decreasing the size of the pore membrane, may help capture more eDNA for sequencing^26^. Regardless of these challenges, we were still able to capture a wide array of target taxonomic groups across the animal tree of life simultaneously, including benthic species that would otherwise each be costly in the form of taxonomic expertise, specialized techniques, and effort to observe through traditional visual means.

Our study supports the feasibility and utility of using molecular-based approaches for quantifying tropical marine biodiversity with relatively low sampling effort. We found the DNA signature of many invertebrate species and, more surprisingly, of numerous fishes that were not observed during visual surveys or never documented in the area. As we hypothesized, eDNA effectively identified patterns of diversity in fish and invertebrate communities in this heterogeneous seascape despite a high level of variability in the dataset likely due to water circulation. We were able to identify environmental features (i.e, exposure) that drive the distribution of animal communities, and we identified sets of habitat specialized species that are consistent with our visual observations. Utilizing environmental DNA sequencing techniques in biodiversity assessments will be crucial as people continue to balance marine resource use and conservation in tropical coastal ecosystems.

## Acknowledgements

We thank the Smithsonian Tropical Research Institute staff at the Bocas del Toro and Naos stations, and the Smithsonian Laboratory of Analytical Biology for their support during this project. We thank William Wied and Daniella Heflin for conducting fish surveys. Sample collection and processing was funded through a Wagoner Foreign Study Scholarship awarded to E.W.S. by Rice University. This study was also supported by a Scholarly Studies Award from the Office of the Provost of the Smithsonian Institution to M.L., A.H.A. and N.K., the Smithsonian Sant Chair for Marine Science and the Global Genome Initiative under the GGI Rolling 2016 and GGI Rolling 2017 Awards Programs to M.L. Sequencing was performed by the George Washington University’s Genomics Core. Computationally-intensive analyses were completed using both the George Washington University’s Colonial One and the Smithsonian Institution’s Hydra high-performance computing systems.

## Author Contributions

E.W.S., B.N.N., J.S. and M.L. designed the study with input from all other co-authors. E.W.S collected the water samples. E.W.S and M.L. extracted DNA from samples and prepared them for sequencing. J.S. led the diver surveys. B.N.N., E.W.S., and M.L. analyzed the data. B.N.N., E.W.S, and M.L. wrote the original draft of the manuscript and all authors reviewed and edited the final version of the manuscript.

## References

1. Costanza, R. et al. Changes in the global value of ecosystem services. Glob. Environ. Chang. 26, 152–158 (2014).

2. Ryther, J. H. Photosynthesis and Fish Production in the Sea. Science (80-.). 166, 72–76 (1969).

3. Millenium Ecosystem Assessment. Ecosystems and Human Well-being: Current State and Trends, Volume 1. Millennium Ecosystem Assessment Series (2005).

4. Gray, J. S. Marine biodiversity: patterns, threats and conservation needs. Biodivers. Conserv. 6, 153–175 (1997).

5. Spalding, M. D., Ravilious, C. & Green, E. P. World Atlas of Coral Reefs. (University of California Press, 2001).

6. Worm, B. & Lenihan, H. S. Threats to Marine Ecosystems. in Marine Community Ecology and Conservation (eds. Bertness, M. D., Bruno, J. F., Silliman, B. R. & Stachowicz, J. J.) 449–476 (Sinauer Associates Inc., 2014).

7. Lotze, H. K. et al. Depletion, Degredation, and Recovery Potential of Estuaries and Coastal Seas. Science (80-.). 312, 1806–1809 (2006).

8. Butchart, S. H. M. et al. Global Biodiversity: Indicators of Recent Declines. Science (80-.). 328, 1164–1168 (2010).

9. Stachowicz, J. J., Bruno, J. F. & Duffy, J. E. Understanding the Effects of Marine Biodiversity on Communities and Ecosystems. Annu. Rev. Ecol. Evol. Syst. 38, 739–766 (2007).

10. Worm, B. et al. Impacts of Biodiversity Loss on Ocean Ecosystem Services. Science (80-.). 314, 787–790 (2006).

11. Duffy, J. E. Reefs need richness. Nat. Ecol. Evol. 3, 149–150 (2019).

12. Topor, Z. M., Rasher, D. B., Duffy, J. E. & Brandl, S. J. Marine protected areas enhance coral reef functioning by promoting fish biodiversity. Conserv. Lett. 12, e12638 (2019).

13. Miloslavich, P. et al. Marine biodiversity in the caribbean: Regional estimates and distribution patterns. PLoS One 5, (2010).

14. Ackerman, J. & Bellwood, D. Reef fish assemblages: a re-evaluation using enclosed rotenone stations. Mar. Ecol. Prog. Ser. 206, 227–237 (2000).

15. Ficetola, G. F., Miaud, C., Pompanon, F. & Taberlet, P. Species detection using environmental DNA from water samples. Biol. Lett. 4, 423–425 (2008).

16. Thomsen, P. F. et al. Detection of a Diverse Marine Fish Fauna Using Environmental DNA from Seawater Samples. PLoS One 7, 1–9 (2012).

17. Thompson, A. & Mapstone, B. Observer effects and training in underwater visual surveys of reef fishes. Mar. Ecol. Prog. Ser. 154, 53–63 (1997).

18. Ackerman, J. L. & Bellwood, D. R. The contribution of small individuals to density-body size relationships. Oecologia 136, 137–140 (2003).

19. Jørgensen, L. L., Renaud, P. E. & Cochrane, S. K. J. Improving benthic monitoring by combining trawl and grab surveys. Mar. Pollut. Bull. 62, 1183–1190 (2011).

20. Boussarie, G., Teichert, N., Lagarde, R. & Ponton, D. BichiCAM, an Underwater Automated Video Tracking System for the Study of Migratory Dynamics of Benthic Diadromous Species in Streams. River Res. Appl. 32, 1392–1401 (2016).

21. Foote, A. D. et al. Investigating the Potential Use of Environmental DNA (eDNA) for Genetic Monitoring of Marine Mammals. PLoS One 7, e41781 (2012).

22. Port, J. A. et al. Assessing vertebrate biodiversity in a kelp forest ecosystem using environmental DNA. Mol. Ecol. 25, 527–541 (2016).

23. Andruszkiewicz, E. A. et al. Biomonitoring of marine vertebrates in Monterey Bay using eDNA metabarcoding. PLoS One 12, e0176343 (2017).

24. DiBattista, J. D. et al. Assessing the utility of eDNA as a tool to survey reef-fish communities in the Red Sea. Coral Reefs 36, 1245–1252 (2017).

25. Nichols, P. K. & Marko, P. B. Rapid assessment of coral cover from environmental DNA in Hawai’i. Environ. DNA 1, 40–53 (2019).

26. Deiner, K. et al. Environmental DNA metabarcoding: Transforming how we survey animal and plant communities. Mol. Ecol. 26, 5872–5895 (2017).

27. O’Donnell, J. L. et al. Spatial distribution of environmental DNA in a nearshore marine habitat. PeerJ 5, e3044 (2017).

28. Stat, M. et al. Ecosystem biomonitoring with eDNA: Metabarcoding across the tree of life in a tropical marine environment. Sci. Rep. 7, 1–11 (2017).

29. Jørgensen, P. S., Folke, C. & Carroll, S. P. Evolution in the Anthropocene: Informing Governance and Policy. Annu. Rev. Ecol. Evol. Syst. 50, annurev-ecolsys-110218-024621 (2019).

30. Edgar, G. J. & Stuart-Smith, R. D. Systematic global assessment of reef fish communities by the Reef Life Survey program. Sci. Data 1, 140007 (2014).

31. Nagelkerken, I. et al. Importance of mangroves, seagrass beds and the shallow coral reef as a nursery for important coral reef fishes, using a visual census technique. Estuar. Coast. Shelf Sci. 51, 31–44 (2000).

32. Nagelkerken, I. et al. How important are mangroves and seagrass beds for coral-reef fish? The nursery hypothesis tested on an island scale. Mar. Ecol. Prog. Ser. 244, 299–305 (2002).

33. Lefcheck, J. S. et al. Are coastal habitats important nurseries? A meta-analysis. Conserv. Lett. e12645 (2019). doi:10.1111/conl.12645

34. Unsworth, R. K. F., Nordlund, L. M. & Cullen-Unsworth, L. C. Seagrass meadows support global fisheries production. Conserv. Lett. 12, 1–8 (2019).

35. Gillis, L. G. Connectivity beyond biodiversity: are physical fluxes important in the tropical coastal seascape? (Radboud University, 2014).

36. Collin, R. Ecological monitoring and biodiversity surveys at the Smithsonian Tropical Research Institute’s Bocas del Toro Research Station. Caribb. J. Sci. 41, 367–373 (2005).

37. Guzmán, H. M., Barnes, P. A. G., Lovelock, C. E. & Feller, I. C. A site description of the CARICOMP mangrove, seagrass and coral reef sites in Bocas del Toro, Panama. Caribb. J. Sci. 41, 430–440 (2005).

38. Dominici-Arosemena, A. & Wolff, M. Reef fish community structure in Bocas del Toro (Caribbean, Panama): Gradients in habitat complexity and exposure. Caribb. J. Sci. 41, 613–637 (2005).

39. De Grave, S. & Anker, A. An annotated checklist of marine caridean and stenopodidean shrimps (Malacostraca: Decapoda) of the Caribbean coast of Panama. Nauplius 25, (2017).

40. Paulay, G. et al. Cryptobenthic invertebrates of Bocas del Toro, Panama: field guide 1.0. (2017). doi:10.6084/m9.figshare.5183722.v1

41. Guzman, H. M. Caribbean coral reefs of Panama: present status and future perspectives. in Latin American coral reefs 241–274 (Elsevier, 2003).

42. Cramer, K. L. et al. Anthropogenic mortality on coral reefs in Caribbean Panama predates coral disease and bleaching. Ecol. Lett. 15, 561–567 (2012).

43. Seemann, J. et al. Assessing the ecological effects of human impacts on coral reefs in Bocas del Toro, Panama. 1747–1763 (2014). doi:10.1007/s10661-013-3490-y

44. Nelson, H. R., Kuempel, C. D. & Altieri, A. H. The resilience of reef invertebrate biodiversity to coral mortality. Ecosphere 7, (2016).

45. Altieri, A. H. et al. Tropical dead zones and mass mortalities on coral reefs. Proc. Natl. Acad. Sci. 114, 3660–3665 (2017).

46. Laramie, M. B., Pilliod, D. S., Goldberg, C. S. & Strickler, K. M. Environmental DNA sampling protocol – filtering water to capture DNA from aquatic organisms. U.S Geol. Surv. Tech. Methods Book 2, 15 p. (2015).

47. Leray, M. et al. A new versatile primer set targeting a short fragment of the mitochondrial COI region for metabarcoding metazoan diversity: Application for characterizing coral reef fish gut contents. Front. Zool. 10, 1–14 (2013).

48. Leray, M., Haenel, Q. & Bourlat, S. J. Preparation of Amplicon Libraries for Metabarcoding of Marine Eukaryotes Using Illumina MiSeq: The Adapter Ligation Method. in Methods in Molecular Biology 1452, 209–218 (Humana Press Inc., 2016).

49. Roehr, J. T., Dieterich, C. & Reinert, K. Flexbar 3.0 – SIMD and multicore parallelization. Bioinformatics 33, 2941–2942 (2017).

50. Callahan, B. J. et al. DADA2: High-resolution sample inference from Illumina amplicon data. Nat. Methods 13, 581–583 (2016).

51. Callahan, B. J., McMurdie, P. J. & Holmes, S. P. Exact sequence variants should replace operational taxonomic units in marker-gene data analysis. ISME J. 11, 2639–2643 (2017).

52. Rognes, T., Flouri, T., Nichols, B., Quince, C. & Mahé, F. VSEARCH: a versatile open source tool for metagenomics. PeerJ (2016). doi:10.7717/peerj.2584

53. Frøslev, T. G. et al. Algorithm for post-clustering curation of DNA amplicon data yields reliable biodiversity estimates. Nat. Commun. 8, (2017).

54. Gao, X., Lin, H., Revanna, K. & Dong, Q. A Bayesian taxonomic classification method for 16S rRNA gene sequences with improved species-level accuracy. BMC Bioinformatics 18, 1–10 (2017).

55. Machida, R. J., Leray, M., Ho, S. L. & Knowlton, N. Data Descriptor: Metazoan mitochondrial gene sequence reference datasets for taxonomic assignment of environmental samples. Sci. Data 4, 1–7 (2017).

56. Humann, P. & DeLoach, N. Reef fish identification: Florida, Caribbean, Bahamas. (New World Publications, 2014).

57. Froese, R. & Pauly, D. FishBase.

58. Robertson, D. R. & Van Tassel, J. Shorefishes of the Greater Caribbean: online information system. Version 2. (2016).

59. R Development Core Team. R: A Language and Environment for Statistical Computing. 1, (R Foundation for Statistical Computing, 2008).

60. McMurdie, P. J. & Holmes, S. phyloseq: An R Package for Reproducible Interactive Analysis and Graphics of Microbiome Census Data. PLoS One 8, e61217 (2013).

61. Oksanen, J. et al. vegan: Community Ecology Package. (2018).

62. Love, M. I., Huber, W. & Anders, S. Moderated estimation of fold change and dispersion for RNA-seq data with DESeq2. Genome Biol. 15, 550 (2014).

63. Aiello, S., Eckstrand, E., Fu, A., Landry, M. & Aboyoun, P. Fast Scalable R with H2O. (2015).

64. Plotly Technologies Inc. Collaborative data science. (2015).

65. Wickham, H. ggplot2: Elegant Graphics for Data Analysis. (Springer-Verlag New York, 2009).

66. Anderson, M. J. Permutational Multivariate Analysis of Variance (PERMANOVA). in Wiley StatsRef: Statistics Reference Online 1–15 (John Wiley & Sons, Ltd, 2017). doi:10.1002/9781118445112.stat07841

67. Woodley, J. D. et al. Hurricane Allen’s Impact on Jamaican Coral Reefs. Science 214, 749–755 (1981).

68. Guzmán, H. M. & Guevara, C. A. Arrecifes coralinos de Bocas del Toro, Panamá: distribución, estructura y estado de conservación de los arrecifes continentales de la Laguna de Chiriquí y la Bahía Almirante. Revista de Biología Tropical 46, 601–623 (1998).

69. Guzmán, H. M. & Guevara, C. A. Arrecifes coralinos de Bocas del Toro, Panamá: II. Distribución, estructura y estado de conservación de los arrecifes de las Islas Bastimentos, Solarte, Carenero y Colón. Revista de Biología Tropical 46, 889–912 (1998).

70. Guzmán, H. M. & Guevara, C. A. Arrecifes coralinos de Bocas del Toro, Panamá: III. Distribución, estructura, diversidad y estado de conservación de los arrecifes de las islas Pastores, Cristóbal, Popa y Cayo Agua. Revista de Biología Tropical 47, 659–676 (1999).

71. Coissac, E., Hollingsworth, P. M., Lavergne, S. & Taberlet, P. From barcodes to genomes: Extending the concept of DNA barcoding. Mol. Ecol. 25, 1423–1428 (2016).

72. Ratnasingham, S. & Hebert, P. D. N. BOLD: The Barcode of Life Data System: Barcoding. Mol. Ecol. Notes 7, 355–364 (2007).

73. Hinchliff, C. E. et al. Synthesis of phylogeny and taxonomy into a comprehensive tree of life. Proc. Natl. Acad. Sci. U. S. A. 112, 12764–12769 (2015).

74. Leray, M. & Knowlton, N. DNA barcoding and metabarcoding of standardized samples reveal patterns of marine benthic diversity. Proc. Natl. Acad. Sci. 2014, 201424997 (2015).

75. Pearman, J. K. et al. Cross-shelf investigation of coral reef cryptic benthic organisms reveals diversity patterns of the hidden majority. Sci. Rep. 8, (2018).

76. Collins, R. A. et al. Non-specific amplification compromises environmental DNA metabarcoding with COI. Methods Ecol. Evol. (2019). doi:10.1111/2041-210x.13276

77. Huang, D., Meier, R., Todd, P. A. & Chou, L. M. Slow mitochondrial COI sequence evolution at the base of the metazoan tree and its implications for DNA barcoding. J. Mol. Evol. 66, 167–174 (2008).

78. Seemann, J., Yingst, A., Stuart-Smith, R. D., Edgar, G. J. & Altieri, A. H. The importance of sponges and mangroves in supporting fish communities on degraded coral reefs in Caribbean Panama. PeerJ 6, e4455 (2018).

