## Supplementary Information for "Environmental DNA survey captures patterns of fish and invertebrate diversity across a tropical seascape"

^1^Computational Biology Institute, The Milken Institute School of Public Health, The George Washington University, Washington, DC, USA. ^2^Department of Biological Sciences, The George Washington University, Washington, DC, USA. ^3^Department of Biological Sciences, University of Rhode Island, Kingston, RI, USA. ^4^Smithsonian Tropical Research Institute, Smithsonian Institution, Panama City, Balboa Ancon, Panama. ^5^Department of Biosciences, Rice University, Houston, TX, USA. ^6^ School of Marine and Environmental Affairs, University of Washington, Seattle, WA, USA. ^7^ Department of Environmental Engineering Sciences, University of Florida, Gainesville, FL, USA. ^8^National Museum of Natural History, Smithsonian Institution, Washington, DC, USA. ^9^Department of Epidemiology and Biostatistics, The Milken Institute School of Public Health, The George Washington University, Washington, DC, USA.

### Supplementary Information

#### Text

##### Fish-specific primer study

A 163-185 bp fragment of the hypervariable region of the 12s rRNA gene (Yamamoto et al. 2016) was amplified using two-step PCR for each sample. The first-stage PCR used two sets of PCR primers designed to target fish and elasmobranchs respectively (Table S1). We modified the protocol as described in Yamamoto et al. (2016) by adding a phosphorothioate bond between the last two bases on the primers to prevent degradation of the primers by the strong proofreading activity by the KAPA HiFi Hotstart ReadyMix. Triplicate 10 µL PCR reactions contained 6 µL of KAPA HiFi HotStart ReadyMix (2x), 0.7 µL of each primer (four primers, each at 5µM), 1.2 µL of molecular-grade water, and 2 µL of DNA extract. The PCRs were run using the following cycling conditions: 3 minutes at 95℃ (1x); 20 s at 95°C, 15 s at 65°C, and 15 s at 72°C (38x); 5 min at 72°C (1x). PCR amplification success was checked using gel electrophoresis on a 3.0% agarose gel. PCR triplicates were pooled, purified using magnetic KAPA Pure Beads (bead:sample ratio of 1.5:1) and quantified using Qubit dsDNA HS assay.

Pooled triplicate PCR products were diluted tenfold with nuclease-free molecular-grade water and used as the template for a second-step PCR reaction to create sequencing-ready Illumina libraries. We substituted the second-step PCR primers described by Yamamoto et al. (2016) in favor of the iTru5 and iTru7 primers described by Glenn et al. (2016), which are identical, except that the Glenn et al. (2016) iTru5 and iTru7 primers have a shorter overlap length (15 bp vs 34 bp) with the product from the first step PCR. Second-step 10 µL PCR reactions contained 6 µL of KAPA HiFi Hotstart ReadyMix (2x), 0.7 µL each of iTru5 and iTru7 primers (each at 0.5 mM), 3.6 µL of molecular-grade water, and 1 µL of template. The PCRs were run using the following cycling conditions: 3 minutes at 95℃ (1x); 30 s at 95°C, 30 s at 60°C, and 30 s at 72°C (12x); 5 min at 72°C (1x). PCR amplification success was checked using gel electrophoresis on a 3.0% agarose gel. PCR products were cleaned using KAPA Pure Beads (bead:sample ratio of 0.9:1) to produce ready-to-sequence libraries. Libraries were quantified using the Qubit dsDNA HS assay.

We encountered little success with the first-stage PCR amplification of the 12S rRNA fragment targeted by the MiFish primers. Out of 142 DNA extracts, only 27 samples had a medium or strong band after gel electrophoresis. The remaining reactions resulted in no bands (n = 42) or very faint bands (n = 73). The pattern of amplification success was consistent across PCR triplicates. Attempts to amplify the first-stage product with the second-stage PCR were also limited in success: most samples did not successfully amplify and multiple bands were observed in samples that did amplify.

#### Figures

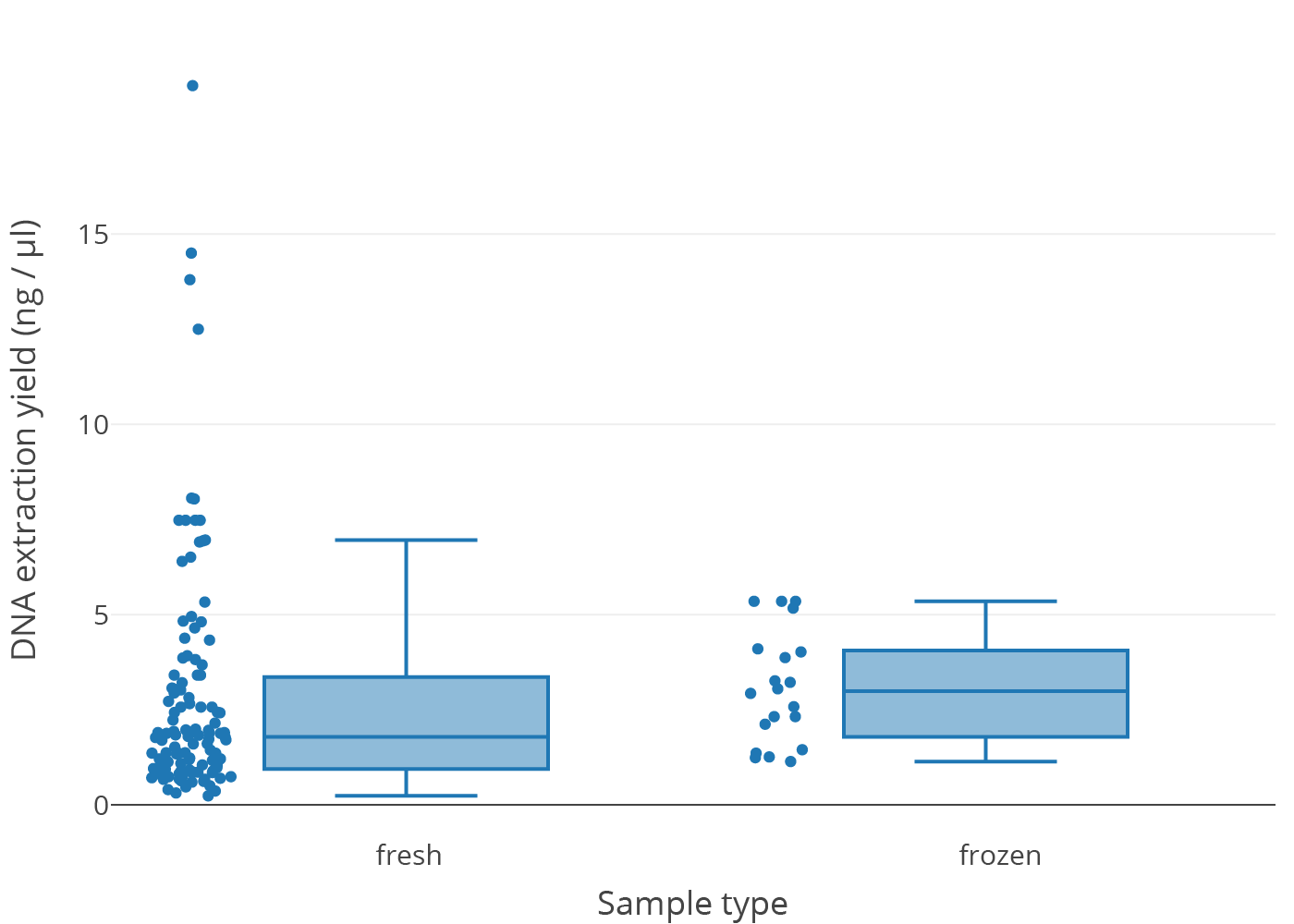

**Supplementary Figure S1. The distribution of DNA extraction yield of immediately filtered samples and frozen samples.** The points show individual samples.

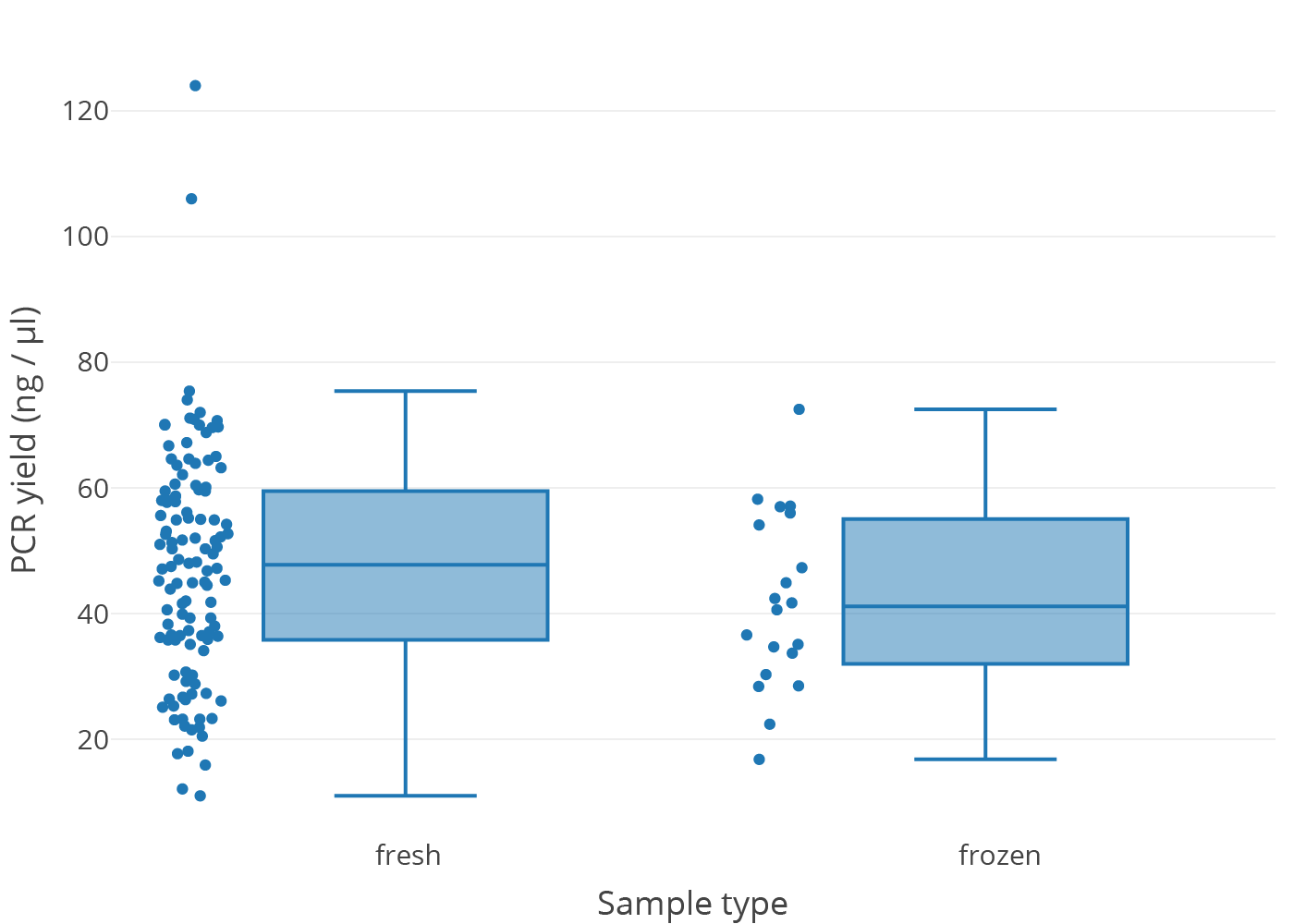

**Supplementary Figure S2. The distribution of PCR yields of immediately filtered samples and frozen samples.** The points show individual samples.

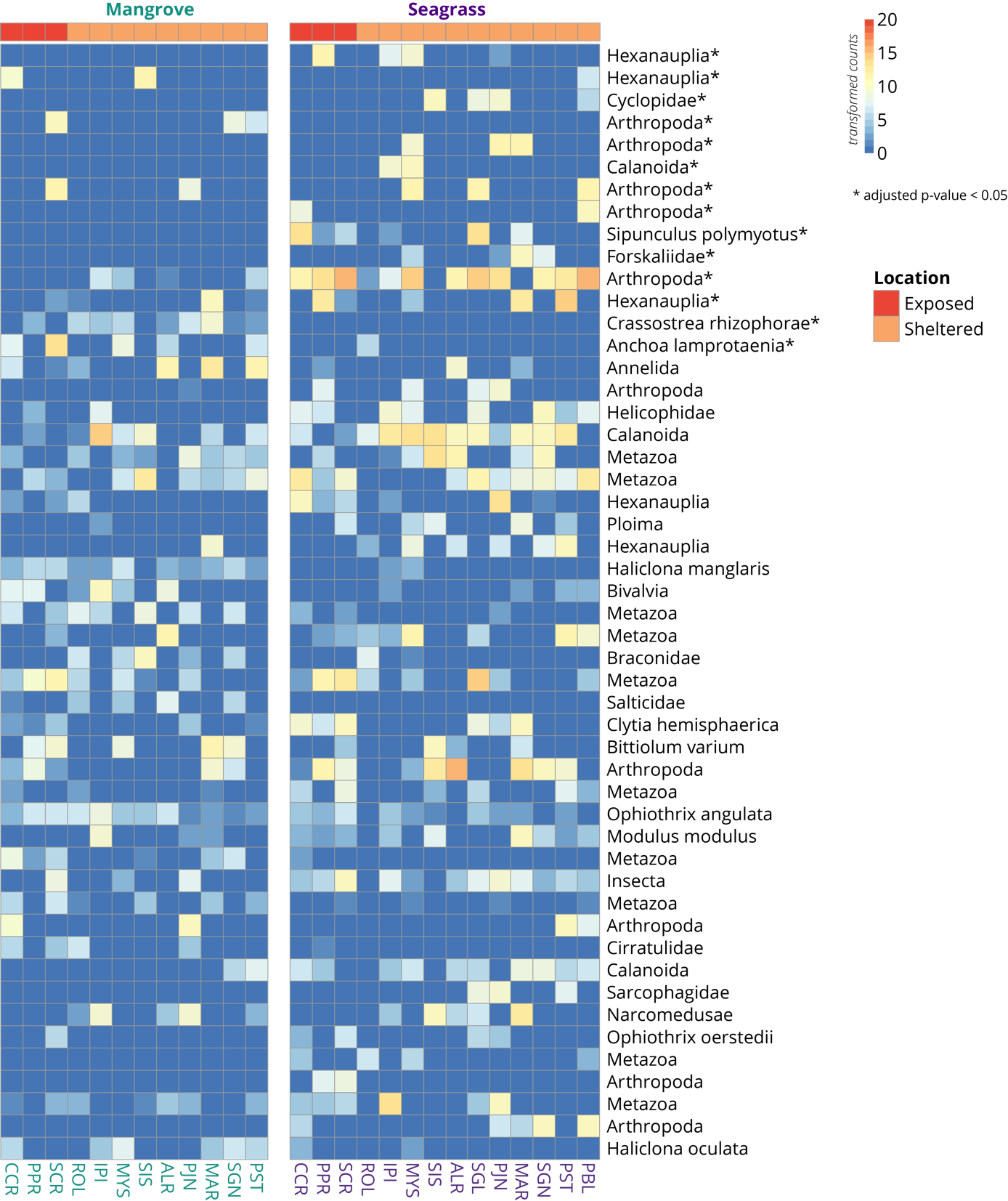

**Supplementary Figure S3. Heatmaps of metazoan taxa differentially abundant in mangrove and seagrass sites**. The 50 metazoan taxa are ordered from top to bottom by ascending adjusted p-values (decreasing significance), with asterisks (*) denoting an adjusted p-value < 0.05. The scale bar and cell colors show count values after variance stabilizing transformation.

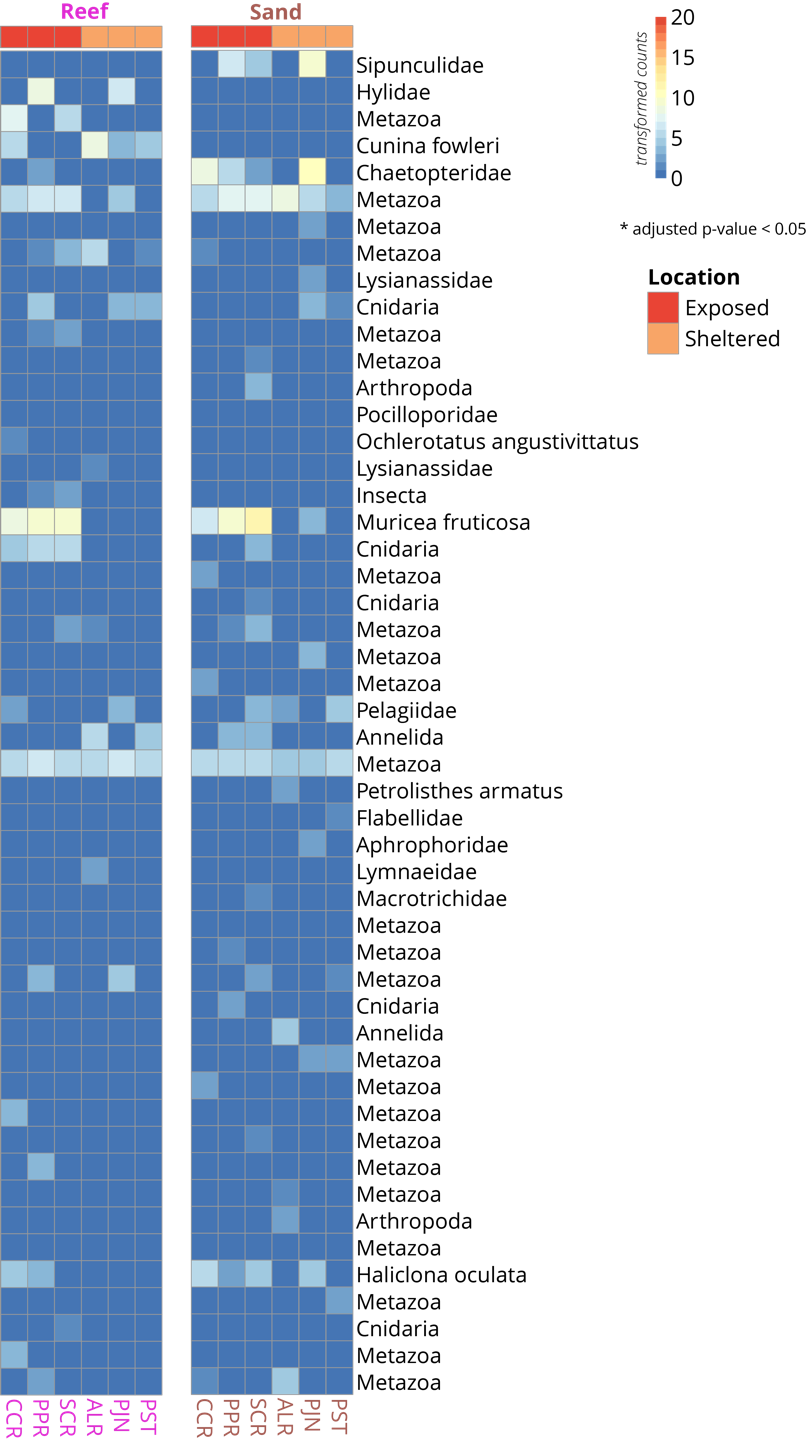

**Supplementary Figure S4. Heatmaps of metazoan taxa differentially abundant in reef and sand sites**. Taxa are ordered top to bottom by ascending adjusted p-values (decreasing significance). No taxa counts were significant (adjusted p-value < 0.05). The scale bar and cell colors show count values after variance stabilizing transformation.

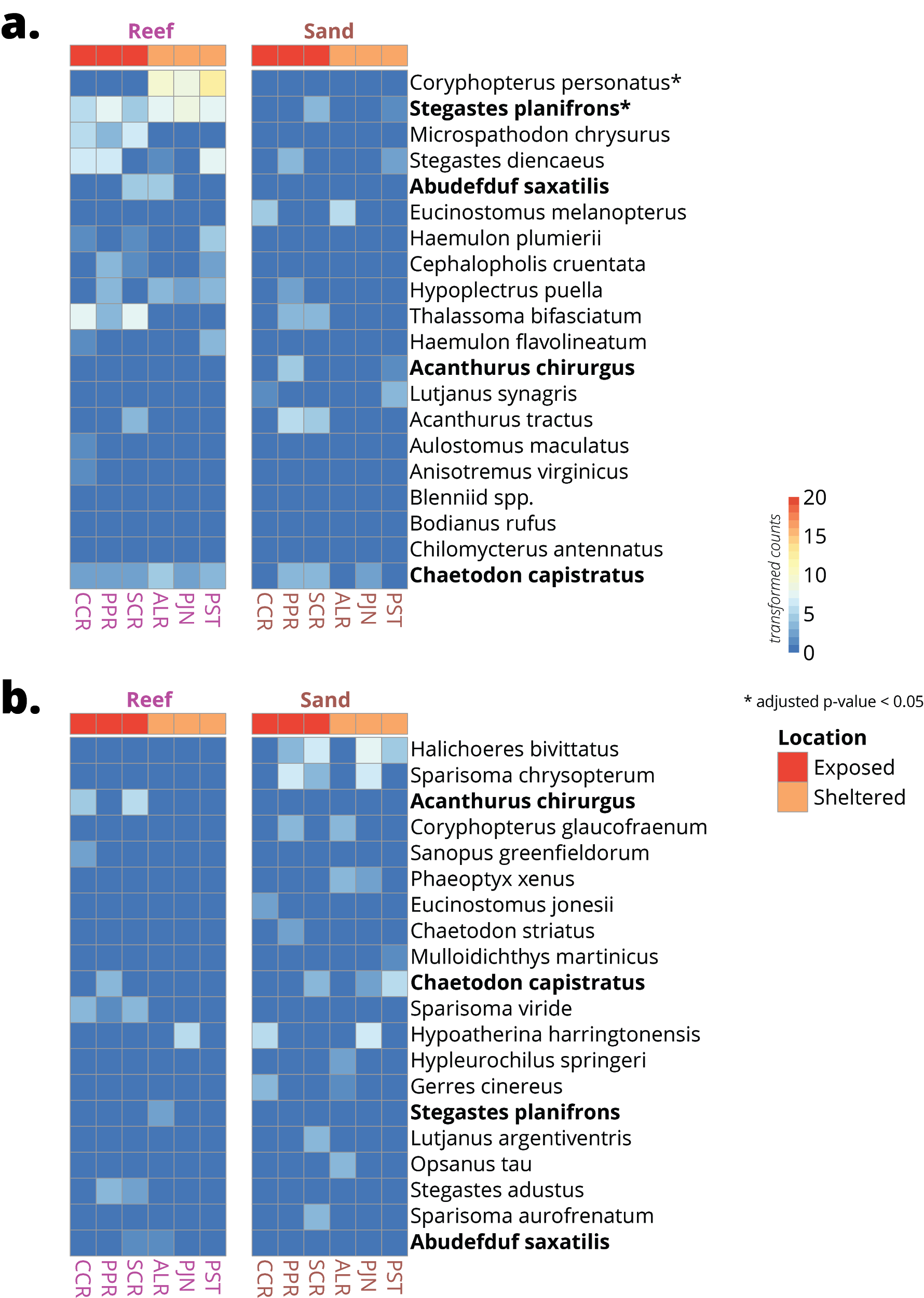

**Supplementary Figure S5. Heatmaps of top 20 differentially detected bony fish taxa using (a) visual surveys and (b) eDNA between coral and sand ecosystems**. The 20 fish taxa are ordered from top to bottom by ascending adjusted p-values (decreasing significance), with asterisks (*) denoting an adjusted p-value < 0.05. Bolded taxa are shared between visual surveys and eDNA. The scale bar and cell colors show count values after variance stabilizing transformation.

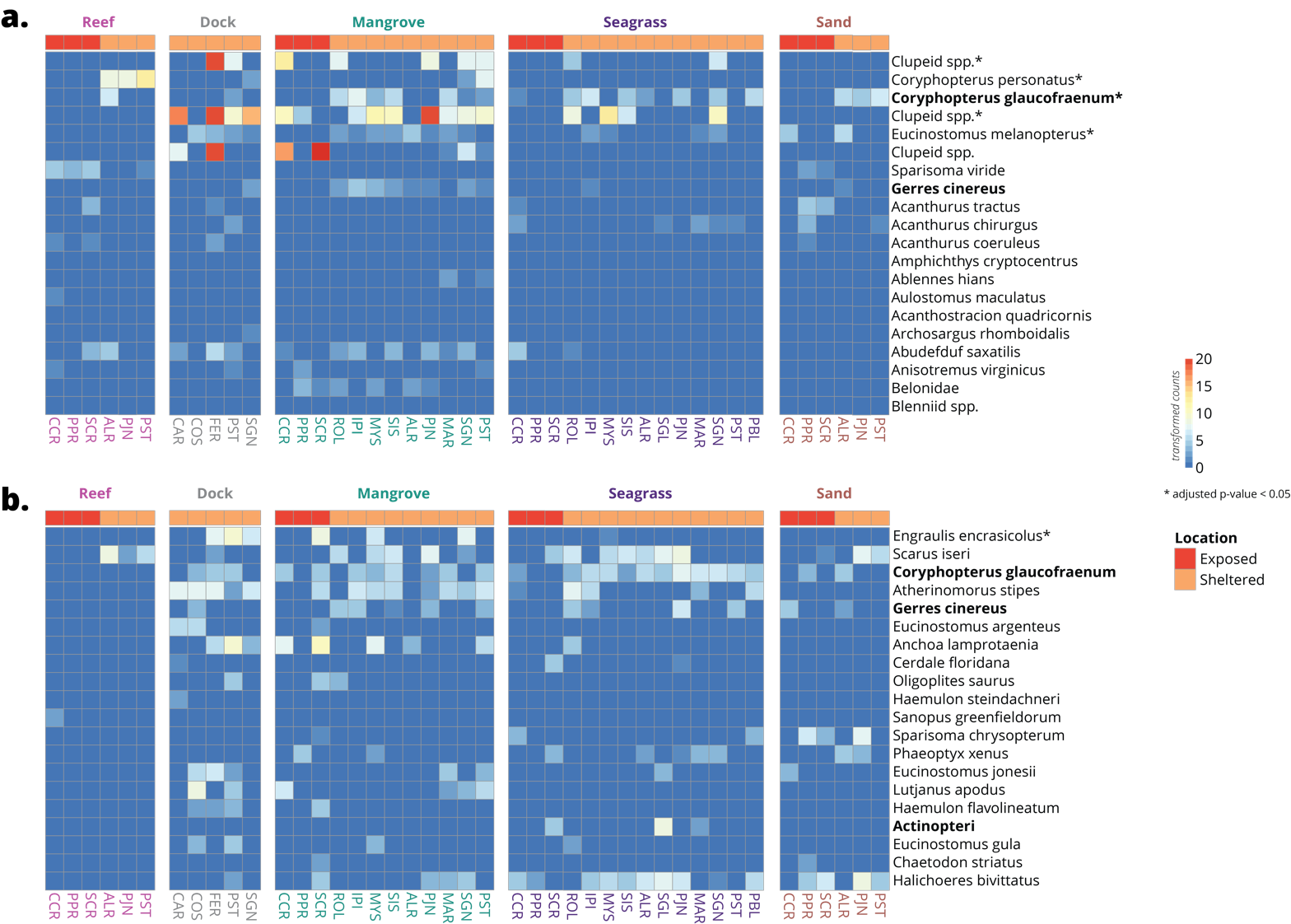

**Supplementary Figure S6. Heatmaps of the top 20 bony fish taxa differentially detected between exposed and sheltered sites in (a) visual surveys and (b) eDNA.** The 20 fish taxa are ordered from top to bottom by ascending adjusted p-values (decreasing significance), with asterisks (*) denoting an adjusted p-value < 0.05. Bolded taxa are shared between visual surveys and eDNA. The scale bar and cell colors show count values after variance stabilizing transformation.

#### Tables

**Supplementary Table S1.** Full site and sampling information. Asterisks (*) denote MarineGeo sites

| Site Name | | Site Code | Latitude | Longitude | Visual survey | eDNA | # Water Samples |
| --- | --- | --- | --- | --- | --- | --- | --- |
| Almirante* | | ALR |  |  |  |  |  |
|  | Coral reef | ALR-C | 9.28993 | -82.34309 | x | x | 3 |
|  | Mangrove | ALR-M | 9.295769 | -82.345434 | x | x | 3 |
|  | Sand | ALR-Sa | 9.295675 | -82.345643 | x | x | 3 |
|  | Seagrass | ALR-S | 9.290707 | -82.342775 | x | x | 3 |
| Coral Cay* | | CCR |  |  |  |  |  |
|  | Coral reef | CCR-C | 9.265417 | -82.119056 | x | x | 3 |
|  | Mangrove | CCR-M | 9.274934 | -82.130763 | x | x | 3 |
|  | Sand | CCR-Sa | 9.274917 | -82.127861 | x | x | 3 |
|  | Seagrass | CCR-S | 9.274167 | -82.127972 | x | x | 3 |
| Pastores Island | | IPI |  |  |  |  |  |
|  | Mangrove | IPI-M | 9.23466 | -82.34544 | x | x | 3 |
|  | Seagrass | IPI-S | 9.23492 | -82.34573 | x | x | 3 |
| Marina | | MAR |  |  |  |  |  |
|  | Mangrove | MAR-M | 9.334212 | -82.247999 | x | x | 3 |
|  | Seagrass | MAR-S | 9.333865 | -82.247656 | x | x | 3 |
| Mystery Spot | | MYS |  |  |  |  |  |
|  | Mangrove | MYS-M | 9.271099 | -82.296867 | x | x | 3 |
|  | Seagrass | MYS-S | 9.2711 | -82.29763 | x | x | 3 |
| Puebla Point | | PBL |  |  |  |  |  |
|  | Seagrass | PBL-S | 9.372683 | -82.291065 | x | x | 3 |
| Juan Point* | | PJN |  |  |  |  |  |
|  | Coral reef | PJN-C | 9.3015 | -82.29404 | x | x | 3 |
|  | Mangrove | PJN-M | 9.2967 | -82.29319 | x | x | 3 |
|  | Sand | PJN-Sa | 9.29728 | -82.29531 | x | x | 3 |
|  | Seagrass | PJN-S | 9.29657 | -82.29321 | x | x | 3 |
| Popa Reef* | | PPR |  |  |  |  |  |
|  | Coral reef | PPR-C | 9.23392 | -82.11185 | x | x | 3 |
|  | Mangrove | PPR-M | 9.231444 | -82.121306 | x | x | 3 |
|  | Sand | PPR-Sa | 9.233952 | -82.117477 | x | x | 3 |
|  | Seagrass | PPR-S | 9.23222 | -82.11339 | x | x | 3 |
| STRI Point* | | PST |  |  |  |  |  |
|  | Coral reef | PST-C | 9.34872 | -82.26258 | x | x | 3 |
|  | Mangrove | PST-M | 9.35195 | -82.258983 | x | x | 3 |
|  | Sand | PST-Sa | 9.34857 | -82.26248 | x | x | 3 |
|  | Seagrass | PST-S | 9.35229 | -82.25833 | x | x | 3 |
| Roldan Cay | | ROL |  |  |  |  |  |
|  | Mangrove | ROL-M | 9.21451 | -82.324 | x | x | 3 |
|  | Seagrass | ROL-S | 9.2147 | -82.324 | x | x | 3 |
| Salt Creek* | | SCR |  |  |  |  |  |
|  | Coral reef | SCR-C | 9.27962 | -82.10162 | x | x | 3 |
|  | Mangrove | SCR-M | 9.28842 | -82.1121 | x | x | 3 |
|  | Sand | SCR-Sa | 9.28293 | -82.10379 | x | x | 3 |
|  | Seagrass | SCR-S | 9.28619 | -82.10524 | x | x | 3 |
| Seagal | | SGL |  |  |  |  |  |
|  | Seagrass | SGL-S | 9.28882 | -82.29497 | x | x | 3 |
| Saigon Bay | | SGN |  |  |  |  |  |
|  | Mangrove | SGN-M | 9.34678 | -82.25704 | x | x | 3 |
|  | Seagrass | SGN-S | 9.34662 | -82.25719 | x | x | 3 |
| Hermanas Cay | | SIS |  |  |  |  |  |
|  | Mangrove | SIS-M | 9.2691 | -82.3517 | x | x | 3 |
|  | Seagrass | SIS-S | 9.268787 | -82.351869 | x | x | 3 |
| Docks | |  |  |  |  |  |  |
|  | Caranero Dock | CAR-D | 9.34217 | -82.23582 | x | x | 2 |
|  | Cosmic Crab Dock | COS-D | 9.343612 | -82.235834 | x | x | 4 |
|  | Ferry Dock | FER-D | 9.335597 | -82.240609 | x | x | 4 |
|  | STRI Dock | PST-D | 9.351037 | -82.257226 | x | x | 2 |
|  | Saigon Dock | SGN-D | 9.34776 | -82.25618 | x | x | 2 |

**Supplementary Table S2.** Details of the primers used in this study. The asterisk represents an added phosphorothioate bond.

| Primer | Sequence | Reference |
| --- | --- | --- |
| **Metazoan** |  |  |
| mlCOIintF | 5’-GGWACWGGWTGAACWGTWTAYCCYCC-3’ | Leray et al. 2013 |
| jgHCO2198 | 5’- TAIACYTCIGGRTGICCRAARAAYCA-3’ | Leray et al. 2013 |
| **MiFish** |  |  |
| MiFish-U-F | 5’-ACACTCTTTCCCTACACGACGCTCTTCCGATCTNNNNNNGTCGGTAAAACTCGTGCCAG*C-3’ | Miya et al. 2015 |
| MiFish-U-R | 5’-GTGACTGGAGTTCAGACGTGTGCTCTTCCGATCTNNNNNNCATAGTGGGGTATCTAATCCCAGTTT*G-3’ | Miya et al. 2015 |
| MiFish-E-F | 5’-ACACTCTTTCCCTACACGACGCTCTTCCGATCTNNNNNNGTTGGTAAATCTCGTGCCAG*C-3’ | Miya et al. 2015 |
| MiFish-E-R | 5’-GTGACTGGAGTTCAGACGTGTGCTCTTCCGATCTNNNNNNCATAGTGGGGTATCTAATCCTAGTTT*G-3’ | Miya et al. 2015 |

**Supplementary Table S3.** Sequencing information

| Sample Name | Full Site Name | Primer Index Name | Primer Index Sequence | Library Index Name | Library Index Sequence |
| --- | --- | --- | --- | --- | --- |
| ROL-M2 | Roldan Cay - Mangrove 2 | Tag_01 | AGACGC | AR002 | CGATGT |
| PPR-Neg | Popa Reef - Neg | Tag_02 | AGTGTA | AR002 | CGATGT |
| SCR-S1 | Salt Creek - Seagrass 1 | Tag_03 | ACTAGC | AR002 | CGATGT |
| IPI-S1 | Pastores Island - Seagrass 1 | Tag_04 | ACAGTC | AR002 | CGATGT |
| PBL-S3 | Puebla Point - Seagrass 3 | Tag_05 | ATCGAC | AR002 | CGATGT |
| MYS-M2 | Mystery Spot - Mangrove 2 | Tag_06 | ATGTCG | AR002 | CGATGT |
| CCR-M2 | Coral Cay - Mangrove 2 | Tag_07 | ATAGCA | AR002 | CGATGT |
| MYS-S1 | Mystery Spot - Seagrass 1 | Tag_08 | AGCTAG | AR002 | CGATGT |
| PJN-C1 | Juan Point - Coral | Tag_09 | ACGTAT | AR002 | CGATGT |
| ALR-Sa3 | Almirante - Sand 3 | Tag_10 | AGTCAT | AR002 | CGATGT |
| SIS-M3 | Hermanas Cay - Mangrove 3 | Tag_11 | AGATCG | AR002 | CGATGT |
| PJN-C2 | Juan Point - Coral | Tag_12 | AGCATC | AR002 | CGATGT |
| MYS-S1 | Mystery Spot - Seagrass 1 | Tag_13 | ACTGAT | AR002 | CGATGT |
| IPI-S2 | Pastores Island - Seagrass 2 | Tag_01 | AGACGC | AR004 | TGACCA |
| PPR-S1 | Popa Reef - Seagrass 1 | Tag_02 | AGTGTA | AR004 | TGACCA |
| ALR-S2 | Almirante - Seagrass 2 | Tag_03 | ACTAGC | AR004 | TGACCA |
| ALR-M1 | Almirante - Mangrove 1 | Tag_04 | ACAGTC | AR004 | TGACCA |
| MAR-S3 | Marina - Seagrass 3 | Tag_05 | ATCGAC | AR004 | TGACCA |
| SGN-Neg | Saigon Bay - Negative | Tag_06 | ATGTCG | AR004 | TGACCA |
| PST-D1 | STRI Point Dock 1 | Tag_07 | ATAGCA | AR004 | TGACCA |
| SCR-M1 | Salt Creek - Mangrove 1 | Tag_08 | AGCTAG | AR004 | TGACCA |
| PPR-Sa3 | Popa Reef - Sand 3 | Tag_09 | ACGTAT | AR004 | TGACCA |
| MAR-M1 | Marina - Mangrove 1 | Tag_10 | AGTCAT | AR004 | TGACCA |
| MAR-M3 | Marina - Mangrove 3 | Tag_11 | AGATCG | AR004 | TGACCA |
| ALR-M3 | Almirante - Mangrove 3 | Tag_12 | AGCATC | AR004 | TGACCA |
| MYS-S1 | Mystery Spot - Seagrass 1 | Tag_13 | ACTGAT | AR004 | TGACCA |
| PJN-C3 | Juan Point - Coral | Tag_01 | AGACGC | AR005 | ACAGTG |
| PPR-M2 | Popa Reef - Mangrove 2 | Tag_02 | AGTGTA | AR005 | ACAGTG |
| SCR-Neg | Salt Creek - Negative | Tag_03 | ACTAGC | AR005 | ACAGTG |
| COS-D1 | Cosmic Crab Dock 1 | Tag_04 | ACAGTC | AR005 | ACAGTG |
| MAR-S2 | Marina - Seagrass 2 | Tag_05 | ATCGAC | AR005 | ACAGTG |
| PST-M3 | STRI Point - Mangrove 3 | Tag_06 | ATGTCG | AR005 | ACAGTG |
| PJN-Sa2 | Juan Point - Sand | Tag_07 | ATAGCA | AR005 | ACAGTG |
| ALR-S1 | Almirante - Seagrass 1 | Tag_08 | AGCTAG | AR005 | ACAGTG |
| PPR-S2 | Popa Reef - Seagrass 2 | Tag_09 | ACGTAT | AR005 | ACAGTG |
| DI-Filter | DI Control (with filter) | Tag_10 | AGTCAT | AR005 | ACAGTG |
| CCR-C3 | Coral Cay - Coral 3 | Tag_11 | AGATCG | AR005 | ACAGTG |
| CCR-Sa1 | Coral Cay - Sand 1 | Tag_12 | AGCATC | AR005 | ACAGTG |
| PJN-C1 | Juan Point - Coral | Tag_13 | ACTGAT | AR005 | ACAGTG |
| PJN-Neg | Juan Point - Negative | Tag_01 | AGACGC | AR006 | GCCAAT |
| SGN-D1 | Saigon Bay Dock 1 | Tag_02 | AGTGTA | AR006 | GCCAAT |
| SGN-S2 | Saigon Bay - Seagrass 2 | Tag_03 | ACTAGC | AR006 | GCCAAT |
| Tap-Neg | Tap water negative | Tag_04 | ACAGTC | AR006 | GCCAAT |
| PJN-S3 | Juan Point - Seagrass | Tag_05 | ATCGAC | AR006 | GCCAAT |
| PJN-Sa3 | Juan Point - Sand | Tag_06 | ATGTCG | AR006 | GCCAAT |
| COS-D2 | Cosmic Crab Dock 2 | Tag_07 | ATAGCA | AR006 | GCCAAT |
| IPI-M1 | Pastores Island - Mangrove 1 | Tag_08 | AGCTAG | AR006 | GCCAAT |
| PST-S2 | STRI Point - Seagrass 2 | Tag_09 | ACGTAT | AR006 | GCCAAT |
| SGN-M1 | Saigon Bay - Mangrove 1 | Tag_10 | AGTCAT | AR006 | GCCAAT |
| PPR-C3 | Popa Reef - Coral 3 | Tag_11 | AGATCG | AR006 | GCCAAT |
| PPR-S3 | Popa Reef - Seagrass 3 | Tag_12 | AGCATC | AR006 | GCCAAT |
| PJN-C1 | Juan Point - Coral | Tag_13 | ACTGAT | AR006 | GCCAAT |
| IPI-M2 | Pastores Island - Mangrove 2 | Tag_01 | AGACGC | AR007 | CAGATC |
| ALR-C2 | Almirante - Coral 2 | Tag_02 | AGTGTA | AR007 | CAGATC |
| CCR-C1 | Coral Cay - Coral 1 | Tag_03 | ACTAGC | AR007 | CAGATC |
| ALR-M2 | Almirante - Mangrove 2 | Tag_04 | ACAGTC | AR007 | CAGATC |
| PJN-S2 | Juan Point - Seagrass | Tag_05 | ATCGAC | AR007 | CAGATC |
| SGN-M3 | Saigon Bay - Mangrove 3 | Tag_06 | ATGTCG | AR007 | CAGATC |
| SCR-C1 | Salt Creek - Coral 1 | Tag_07 | ATAGCA | AR007 | CAGATC |
| DI-Nofilter | DI Control (without filter) | Tag_08 | AGCTAG | AR007 | CAGATC |
| ALR-Sa1 | Almirante - Sand 1 | Tag_09 | ACGTAT | AR007 | CAGATC |
| FER1-D1 | Ferry Dock 1.1 | Tag_10 | AGTCAT | AR007 | CAGATC |
| PST-Sa1 | STRI Point - Sand 1 | Tag_11 | AGATCG | AR007 | CAGATC |
| MYS-S3 | Mystery Spot - Seagrass 3 | Tag_12 | AGCATC | AR007 | CAGATC |
| COS-D1 | Cosmic Crab Dock 1 | Tag_13 | ACTGAT | AR007 | CAGATC |
| PPR-M3 | Popa Reef - Mangrove 3 | Tag_01 | AGACGC | AR012 | CTTGTA |
| PST-Neg | STRI Point - Negative | Tag_02 | AGTGTA | AR012 | CTTGTA |
| IPI-S3 | Pastores Island - Seagrass 3 | Tag_03 | ACTAGC | AR012 | CTTGTA |
| SCR-Sa1 | Salt Creek - Sand 1 | Tag_04 | ACAGTC | AR012 | CTTGTA |
| PJN-Sa1 | Juan Point - Sand | Tag_05 | ATCGAC | AR012 | CTTGTA |
| ROL-S3 | Roldan Cay - Seagrass 3 | Tag_06 | ATGTCG | AR012 | CTTGTA |
| SCR-M3 | Salt Creek - Mangrove 3 | Tag_07 | ATAGCA | AR012 | CTTGTA |
| SGL-S3 | Seagal - Seagrass 3 | Tag_08 | AGCTAG | AR012 | CTTGTA |
| PST-Sa3 | STRI Point - Sand 3 | Tag_09 | ACGTAT | AR012 | CTTGTA |
| ROL-Neg | Roldan Cay - Negative | Tag_10 | AGTCAT | AR012 | CTTGTA |
| PST-M2 | STRI Point - Mangrove 2 | Tag_11 | AGATCG | AR012 | CTTGTA |
| PST-S1 | STRI Point - Seagrass 1 | Tag_12 | AGCATC | AR012 | CTTGTA |
| COS-D1 | Cosmic Crab Dock 1 | Tag_13 | ACTGAT | AR012 | CTTGTA |
| CCR-Sa2 | Coral Cay - Sand 2 | Tag_01 | AGACGC | AR013 | AGTCAA |
| PJN-M2.2 | Juan Point - Mangrove | Tag_02 | AGTGTA | AR013 | AGTCAA |
| CAR-D2 | Caranero Marina Dock 2 | Tag_03 | ACTAGC | AR013 | AGTCAA |
| PPR-Sa2 | Popa Reef - Sand 2 | Tag_04 | ACAGTC | AR013 | AGTCAA |
| SIS-S1 | Hermanas Cay - Seagrass 1 | Tag_05 | ATCGAC | AR013 | AGTCAA |
| SGN-M2 | Saigon Bay - Mangrove 2 | Tag_06 | ATGTCG | AR013 | AGTCAA |
| ALR-C1 | Almirante - Coral 1 | Tag_07 | ATAGCA | AR013 | AGTCAA |
| SIS-Neg | Hermanas Cay - Negative | Tag_08 | AGCTAG | AR013 | AGTCAA |
| PST-C2 | STRI Point - Coral 2 | Tag_09 | ACGTAT | AR013 | AGTCAA |
| SIS-S2 | Hermanas Cay - Seagrass 2 | Tag_10 | AGTCAT | AR013 | AGTCAA |
| PST-D2 | STRI Point Dock 2 | Tag_11 | AGATCG | AR013 | AGTCAA |
| SCR-M2 | Salt Creek - Mangrove 2 | Tag_12 | AGCATC | AR013 | AGTCAA |
| ALR-Sa2 | Almirante - Sand 2 | Tag_13 | ACTGAT | AR013 | AGTCAA |
| PBL-S1 | Puebla Point - Seagrass 1 | Tag_01 | AGACGC | AR014 | AGTTCC |
| FER2-D1 | Ferry Dock 2.1 | Tag_02 | AGTGTA | AR014 | AGTTCC |
| FER1-D2 | Ferry Dock 1.2 | Tag_03 | ACTAGC | AR014 | AGTTCC |
| SIS-M2 | Hermanas Cay - Mangrove 2 | Tag_04 | ACAGTC | AR014 | AGTTCC |
| SGN-D2 | Saigon Bay Dock 2 | Tag_05 | ATCGAC | AR014 | AGTTCC |
| ROL-M3 | Roldan Cay - Mangrove 3 | Tag_06 | ATGTCG | AR014 | AGTTCC |
| CCR-S1 | Coral Cay - Seagrass 1 | Tag_07 | ATAGCA | AR014 | AGTTCC |
| FER2-D2 | Ferry Dock 2.2 | Tag_08 | AGCTAG | AR014 | AGTTCC |
| ALR-Neg | Almirante - Negative | Tag_09 | ACGTAT | AR014 | AGTTCC |
| CCR-M3 | Coral Cay - Mangrove 3 | Tag_10 | AGTCAT | AR014 | AGTTCC |
| PJN-S1 | Juan Point - Seagrass | Tag_11 | AGATCG | AR014 | AGTTCC |
| PPR-M1 | Popa Reef - Mangrove 1 | Tag_12 | AGCATC | AR014 | AGTTCC |
| ALR-Sa2 | Almirante - Sand 2 | Tag_13 | ACTGAT | AR014 | AGTTCC |
| SCR-Sa3 | Salt Creek - Sand 3 | Tag_01 | AGACGC | AR015 | ATGTCA |
| PBL-S2 | Puebla Point - Seagrass 2 | Tag_02 | AGTGTA | AR015 | ATGTCA |
| CCR-C2 | Coral Cay - Coral 2 | Tag_03 | ACTAGC | AR015 | ATGTCA |
| PST-C3 | STRI Point - Coral 3 | Tag_04 | ACAGTC | AR015 | ATGTCA |
| SCR-S2 | Salt Creek - Seagrass 2 | Tag_05 | ATCGAC | AR015 | ATGTCA |
| PST-M1 | STRI Point - Mangrove 1 | Tag_06 | ATGTCG | AR015 | ATGTCA |
| MAR-M2 | Marina - Mangrove 2 | Tag_07 | ATAGCA | AR015 | ATGTCA |
| SCR-S3 | Salt Creek - Seagrass 3 | Tag_08 | AGCTAG | AR015 | ATGTCA |
| SGL-S2 | Seagal - Seagrass 2 | Tag_09 | ACGTAT | AR015 | ATGTCA |
| SIS-S3 | Hermanas Cay - Seagrass 3 | Tag_10 | AGTCAT | AR015 | ATGTCA |
| ROL-M1 | Roldan Cay - Mangrove 1 | Tag_11 | AGATCG | AR015 | ATGTCA |
| MAR-S1 | Marina - Seagrass 1 | Tag_12 | AGCATC | AR015 | ATGTCA |
| Mock | Fort Pierce | Tag_13 | ACTGAT | AR015 | ATGTCA |
| PST-C1.2 | STRI Point - Coral 1 (2/2) | Tag_01 | AGACGC | AR016 | CCGTCC |
| SGN-S3 | Saigon Bay - Seagrass 3 | Tag_02 | AGTGTA | AR016 | CCGTCC |
| CCR-M1 | Coral Cay - Mangrove 1 | Tag_03 | ACTAGC | AR016 | CCGTCC |
| ALR-Sa2 | Almirante - Sand 2 | Tag_04 | ACAGTC | AR016 | CCGTCC |
| SIS-M1 | Hermanas Cay - Mangrove 1 | Tag_05 | ATCGAC | AR016 | CCGTCC |
| PST-Sa2 | STRI Point - Sand 2 | Tag_06 | ATGTCG | AR016 | CCGTCC |
| MYS-M1 | Mystery Spot - Mangrove 1 | Tag_07 | ATAGCA | AR016 | CCGTCC |
| PPR-C2 | Popa Reef - Coral 2 | Tag_08 | AGCTAG | AR016 | CCGTCC |
| ALR-C3 | Almirante - Coral 3 | Tag_09 | ACGTAT | AR016 | CCGTCC |
| CAR-D1 | Caranero Marina Dock 1 | Tag_10 | AGTCAT | AR016 | CCGTCC |
| SGN-S1 | Saigon Bay - Seagrass 1 | Tag_11 | AGATCG | AR016 | CCGTCC |
| PJN-M3 | Juan Point - Mangrove | Tag_12 | AGCATC | AR016 | CCGTCC |
| Mock | Fort Pierce | Tag_13 | ACTGAT | AR016 | CCGTCC |
| CCR-S3 | Coral Cay - Seagrass 3 | Tag_01 | AGACGC | AR018 | GTCCGC |
| SCR-C2 | Salt Creek - Coral 2 | Tag_02 | AGTGTA | AR018 | GTCCGC |
| CCR-Sa3 | Coral Cay - Sand 3 | Tag_03 | ACTAGC | AR018 | GTCCGC |
| 7/12-Neg | 7/12/2017 Negative | Tag_04 | ACAGTC | AR018 | GTCCGC |
| DOCK-Neg | Dock Negative | Tag_05 | ATCGAC | AR018 | GTCCGC |
| IPI-M3 | Pastores Island - Mangrove 3 | Tag_06 | ATGTCG | AR018 | GTCCGC |
| PPR-C1 | Popa Reef - Coral 1 | Tag_07 | ATAGCA | AR018 | GTCCGC |
| CCR-Neg | Coral Cay - Negative | Tag_08 | AGCTAG | AR018 | GTCCGC |
| CCR-S2 | Coral Cay - Seagrass 2 | Tag_09 | ACGTAT | AR018 | GTCCGC |
| MYS-M3 | Mystery Spot - Mangrove 3 | Tag_10 | AGTCAT | AR018 | GTCCGC |
| SCR-C3 | Salt Creek - Coral 3 | Tag_11 | AGATCG | AR018 | GTCCGC |
| PST-M1 | STRI Point - Mangrove 1 | Tag_12 | AGCATC | AR018 | GTCCGC |
| Mock | Fort Pierce | Tag_13 | ACTGAT | AR018 | GTCCGC |
| ALR-S3 | Almirante - Seagrass 3 | Tag_01 | AGACGC | AR019 | GTGAAA |
| SGL-S1 | Seagal - Seagrass 1 | Tag_02 | AGTGTA | AR019 | GTGAAA |
| SCR-Sa2 | Salt Creek - Sand 2 | Tag_03 | ACTAGC | AR019 | GTGAAA |
| ROL-S2 | Roldan Cay - Seagrass 2 | Tag_04 | ACAGTC | AR019 | GTGAAA |
| ROL-S1 | Roldan Cay - Seagrass 1 | Tag_05 | ATCGAC | AR019 | GTGAAA |
| PJN-M1 | Juan Point - Mangrove | Tag_06 | ATGTCG | AR019 | GTGAAA |
| MAR-Neg | Marina - Negative | Tag_07 | ATAGCA | AR019 | GTGAAA |
| IPI-Neg | Pastores Island - Negative | Tag_08 | AGCTAG | AR019 | GTGAAA |
| MYS-S2 | Mystery Spot - Seagrass 2 | Tag_09 | ACGTAT | AR019 | GTGAAA |
| PST-S3 | STRI Point - Seagrass 3 | Tag_10 | AGTCAT | AR019 | GTGAAA |
| PPR-Sa1 | Popa Reef - Sand 1 | Tag_11 | AGATCG | AR019 | GTGAAA |
| PST-M1 | STRI Point - Mangrove 1 | Tag_12 | AGCATC | AR019 | GTGAAA |

**Supplementary Table S4.** Fish species identified during eDNA and visual surveys. * denotes fish species found in the Bocas database, while ^ denotes fish species found in the OBIS database, but not in the Bocas database. † denotes taxonomically misassigned species that were manually checked and corrected. 6 additional fish OTUs in the eDNA were not identified to the genus or species level and not reported here.

| eDNA (43) | eDNA & RLS (36) | RLS (61) |
| --- | --- | --- |
| *Acanthemblemaria aspera* | *Abudefduf saxatilis** | *Ablennes hians* |
| *Acanthurus bahianus** | *Acanthurus chirurgus** | *Acanthostracion quadricornis** |
| *Aetobatus narinari*† | *Carangoides bartholomaei** | *Acanthurus coeruleus** |
| *Anchoa lamprotaenia* | *Caranx crysos** | *Acanthurus tractus* |
| *Anchoa mitchilli* | *Centropomus undecimalis** | *Amphichthys cryptocentrus** |
| *Atherinomorus stipes* | *Chaetodon capistratus** | *Anisotremus virginicus** |
| *Calamus penna** | *Chaetodon striatus** | *Archosargus rhomboidalis** |
| *Caranx sexfasciatus* | *Chloroscombrus chrysurus** | *Aulostomus maculatus** |
| *Cerdale floridana* | *Coryphopterus glaucofraenum^* | *Bodianus rufus* |
| *Coryphopterus dicrus** | *Gerres cinereus** | *Calamus pennatula* |
| *Ctenogobius saepepallens** | *Haemulon aurolineatum** | *Cantherhines pullus** |
| *Diapterus rhombeus^* | *Dasyatis americana** | *Canthigaster rostrata** |
| *Engraulis encrasicolus* | *Haemulon flavolineatum** | *Caranx ruber* |
| *Enneanectes boehlkei* | *Haemulon plumierii** | *Caranx bartholomaei** |
| *Eucinostomus argenteus* | *Haemulon steindachneri^* | *Caranx latus** |
| *Eucinostomus gula^* | *Halichoeres bivittatus** | *Cephalopholis cruentata** |
| *Eucinostomus jonesii* | *Hypleurochilus geminatus* | *Chaetodipterus faber** |
| *Euthynnus alletteratus* | *Hypoplectrus nigricans** | *Chaetodon ocellatus** |
| *Haemulon bonariense^* | *Lophogobius cyprinoides* | *Chilomycterus antennatus* |
| *Hemiramphus brasiliensis^* | *Lutjanus analis** | *Coryphopterus personatus** |
| *Hypanus guttatus* | *Lutjanus apodus** | *Diodon hystrix* |
| *Hypleurochilus springeri* | *Lutjanus griseus** | *Echeneis neucratoides** |
| *Hypoatherina harringtonensis* | *Lutjanus synagris** | *Elacatinus illecebrosus* |
| *Jenkinsia lamprotaenia^* | *Microgobius signatus* | *Eucinostomus melanopterus** |
| *Lutjanus argentiventris* | *Ocyurus chrysurus** | *Gambusia nicaraguensis* |
| *Lutjanus mahogoni** | *Oligoplites saurus^* | *Gnatholepis thompsoni** |
| *Mugil rubrioculus* | *Scarus iseri** | *Haemulon carbonarium** |
| *Mulloidichthys martinicus** | *Serranus tigrinus** | *Haemulon macrostomum** |
| *Opsanus tau* | *Sparisoma aurofrenatum** | *Haemulon parra** |
| *Phaeoptyx pigmentaria* | *Sparisoma chrysopterum** | *Halichoeres maculipinna** |
| *Phaeoptyx xenus^* | *Sparisoma radians** | *Halichoeres pictus** |
| *Poecilia mexicana** | *Sparisoma viride** | *Halichoeres poeyi** |
| *Rhinoptera sp.*† | *Sphyraena barracuda** | *Holacanthus ciliaris* |
| *Rypticus carpenteri* | *Stegastes adustus** | *Hypoplectrus gemma* |
| *Sanopus greenfieldorum* | *Stegastes planifrons** | *Hypoplectrus puella** |
| *Sparisoma rubripinne** | *Urobatis jamaicensis** | *Hypoplectrus unicolor** |
| *Stathmonotus stahli* |  | *Lutjanus cyanopterus** |
| *Strongylura timucu* |  | *Lutjanus jocu** |
| *Thunnus obesus* |  | *Microgobius meeki* |
| *Tomicodon lavettsmithi* |  | *Microgobius microlepis* |
| *Trachinotus falcatus* |  | *Microspathodon chrysurus** |
| *Tylosurus acus* |  | *Monacanthus tuckeri** |
| *Tylosurus crocodilus* |  | *Narcine brasiliensis* |
|  |  | *Odontoscion dentex** |
|  |  | *Ophioblennius atlanticus** |
|  |  | *Opistognathus aurifrons** |
|  |  | *Parablennius marmoreus** |
|  |  | *Paraclinus nigripinnis* |
|  |  | *Pomacanthus arcuatus** |
|  |  | *Prognathodes aculeatus* |
|  |  | *Pseudupeneus maculatus** |
|  |  | *Pterois volitans** |
|  |  | *Scarus trispinosus* |
|  |  | *Scomberomorus maculatus** |
|  |  | *Scomberomorus regalis** |
|  |  | *Serranus tortugarum** |
|  |  | *Sphoeroides spengleri** |
|  |  | *Stegastes diencaeus** |
|  |  | *Stegastes partitus** |
|  |  | *Thalassoma bifasciatum** |
|  |  | *Tigrigobius saucrus* |
